# Comparative single-cell profiling reveals distinct cardiac resident macrophages essential for zebrafish heart regeneration

**DOI:** 10.1101/2022.11.22.517520

**Authors:** Ke-Hsuan Wei, I-Ting Lin, Kaushik Chowdhury, Kuan-Ting Liu, Tai-Ming Ko, Yao-Ming Chang, Kai-Chien Yang, Shih-Lei Lai

## Abstract

Zebrafish exhibit a robust ability to regenerate their hearts following injury, and the immune system plays a key role in this process. We previously showed that delaying macrophage recruitment by clodronate liposome (CL) treatment compromises neutrophil resolution and heart regeneration, even when the infiltrating macrophage number was restored within the first-week post injury (Lai et al., 2017). Here, we examined the molecular mechanisms underlying the cardiac repair of regenerative PBS-control hearts vs. non-regenerative CL-treated hearts. Bulk transcriptomic analyses revealed that CL-treated hearts exhibited disrupted inflammatory resolution and energy metabolism during cardiac repair. Temporal single-cell profiling of inflammatory cells in regenerative vs. non-regenerative conditions further identified heterogenous macrophages and neutrophils with distinct infiltration dynamics, gene expression, and cellular crosstalk. Among them, two residential macrophage subpopulations were enriched in regenerative hearts and barely recovered in non-regenerative hearts. Early CL treatment at 8 days or even 1 month before cryoinjury led to the depletion of resident macrophages without affecting the circulating macrophage recruitment to the injured area. Strikingly, these resident macrophage-deficient zebrafish still exhibited compromised neovascularization and scar resolution. Our results characterized the inflammatory cells of the zebrafish injured hearts and identified key resident macrophage subpopulations prerequisite for successful heart regeneration.

## Introduction

Heart failure is a major cause of morbidity and mortality, in part due to the inability of the human heart to replenish lost cardiomyocytes following cardiac injury such as myocardial infarction (MI). Unlike adult mice and humans, many vertebrates, including certain fish and amphibians, are capable of endogenous heart regeneration throughout life. As an example, zebrafish display a distinct ability to regenerate their heart following injury. However, this ability is not shared by another teleost, the medaka (1, 2). Even though this capacity is lost in adulthood, many mammals can regenerate their heart during embryonic and neonatal stages (3, 4). Comparative studies between neonatal and adult mice (5, 6), and between phylogenetically related species such as zebrafish (*Danio rerio*) and medaka (*Oryzias latipes*) (1, 2) have suggested that the capacity for regeneration does not solely rely on genetic makeup, environmental conditions (e.g., hypoxia), or evolutionary complexity; instead, the type and extent of the immune responses to cardiac injury seem to be a major difference between these regenerative and non-regenerative models (1), and may largely influence the recovery post-experimental MI, as well as clinical prognosis (7).

In our earlier study of reciprocal analyses in zebrafish and medaka, we observed delayed and reduced macrophage recruitment in medaka compared to zebrafish following cardiac injury. Furthermore, delaying macrophage recruitment in zebrafish by intraperitoneal (IP) injection of clodronate liposome (CL) 1 day prior to cryoinjury compromised neovascularization, neutrophil clearance, cardiomyocyte proliferation, and scar resolution, even though the number of infiltrating macrophages recovered to the control levels in the first-week post injury. These results indicate that late cardiac macrophages in CL-treated fish were different in their identity and regenerative potential. We therefore sought to use non-regenerative zebrafish as a comparative model to elucidate the molecular properties of regeneration-associated macrophage. Recent studies in zebrafish identified novel macrophage populations by gene-specific reporters and examined their functions in cardiac repair and regeneration, hinting at a heterogeneous spectrum of cardiac macrophages (8–12). However, beyond these observations, the heterogeneity and function of these inflammatory cells during cardiac regeneration remain unclear.

In this study, we perform comparative bulk and single-cell transcriptomic profiling to investigate the global influence of macrophage pre-depletion and comprehensively analyze the heterogeneity, dynamic, and function of both macrophages and neutrophils in regenerative (PBS-treated) vs. non-regenerative (CL-treated) zebrafish hearts. Bulk RNAseq analysis indicated prolonged and unresolved inflammatory response and mis-regulated energy metabolism in CL-treated zebrafish until 3 weeks post cardiac injury, while cardiomyocyte replenishment and scar resolution took place extensively in regenerative PBS-treated control hearts. Single-cell analyses further revealed diverse macrophage subpopulations with potential functions in phagocytosis, neutrophil recruitment, reactive oxygen species (ROS) homeostasis, angiogenesis, extracellular matrix (ECM) remodeling, and inflammatory regulation during the first-week post cardiac injury. Comparative analyses between regenerative and non-regenerative hearts led to the identification of unique cardiac resident macrophage subpopulations expressing *timp4.3* and *hemoglobin* genes that potentially function in ECM remodeling, inflammatory resolution, and ROS homeostasis. Pre-depleting these resident macrophages a week or even a month prior to cardiac injury significantly impaired heart regeneration without affecting macrophage recruitment from circulation, suggesting that these resident macrophages determine the regenerative capacity and cannot be replaced by circulating/monocyte-derived macrophages. Altogether, these results unravel the heterogeneity and function of inflammatory cells during cardiac repair and highlight the indispensable role of cardiac resident macrophages in zebrafish cardiac regeneration.

## Materials and Methods

### Experimental models

All zebrafish were maintained at our in-house fish facility, Institute of Biomedical Sciences, Academia Sinica using standard husbandry protocol. All experiments were done by institutional and ethical welfare guidelines approved by the ethics committee, Academia Sinica. We crossed *Tg(mpeg1.4:mCherry-F)^ump2^* (13) and *TgBAC(mpx:GFP)^i114^* (14) to generate *Tg(mpx:GFP;mpeg1.4:mCherry-F)* line as well as *Tg(−5.1myl7:DsRed2-NLS)^f2Tg^* (15) and *Tg(fli1:EGFP)* to generate *Tg(fli1:EGFP;-5.1myl7:DsRed2-NLS)* line respectively for our animal experiments. Intraperitoneal (IP) injections of 10 μl PBS and Clodronate liposomes (5 mg/ml) (Liposoma, Amsterdam, The Netherlands) in each fish were performed according to the experimental design.

### Cryoinjury

Cryoinjury was performed as previously described in zebrafish (16–18). In brief, fish were anesthetized in 0.04% tricaine (Sigma, St Louis, MO, USA) and immobilized in a damp sponge with ventral side up. A small incision was created through the thoracic wall by microdissection scissors. A stainless steel cryoprobe precooled in liquid nitrogen was placed on the ventricular surface until thawing could be observed. Fish were placed in a tank of freshwater for recovery, and their reanimation could be enhanced by pipetting water onto the gills after surgery.

### Cryosections and histological analyses

Zebrafish hearts were extracted and fixed in 4% (wt/vol) paraformaldehyde (Alfa Aesar, MA, USA) at room temperature for 1 hour (hr). Collected hearts were subsequently cryopreserved with 30% (wt/vol) sucrose, followed by immersed in OCT (Tissue-Tek, Sakura Finetek, Torrance, CA) and stored at −80°C immediately. 10 μm cryosections were collected for histological analysis. AFOG staining was applied for the visualization of healthy CMs as orange, collagens as blue and fibrins as red. In brief, samples were incubated in preheated Bouin’s solution (Sigma, St Louis, MO, USA) at 58°C for 2 hr post fixation and subsequently immersed in 1% phosphomolybdic acid (Sigma, St Louis, MO, USA) and 0.5% phosphotungstic acid solution (Sigma, St Louis, MO, USA) at room temperature for 5 min for mordanting. Samples were then incubated with AFOG solution containing Aniline Blue (Sigma, St Louis, MO, USA), Orange G (Sigma, St Louis, MO, USA) and Acid Fuchsin (Sigma, St Louis, MO, USA) for development. Quantification for each heart was done by measuring the scar area from the discontinuous sections with the largest scar as well as the two sections at the front and the two sections at the back as previously described (1). Statistic was performed on Prism 9 using student’s t-test.

### Quantitative polymerase chain reaction (qPCR)

For collecting single cells from hearts, 45-60 ventricles were isolated in each experiment at respective time points and conditions, pooled together and digested using LiberaseDH (Sigma, St Louis, MO, USA) at 28C’ in a 24-well plate. Cell suspension was filtered through 100 μm, 70 μm, and 40 μm cell strainers (SPL, Gyeonggi-do, Korea) and centrifuged at 200g for 5 min. Cell pellet was resuspended in 0.04% BSA/PBS, stained with DAPI, and filtered through 35 μm Flowmi cell strainer (BD, New Jersey, USA). Cells were sorted using Fluorescence-activated Cell Sorting (FACS; BD Facs Aria) and then subjected to RNA isolation. RNA was extracted by using TRIzol™ LS Reagent (Life Technologies Invitrogen, CA, USA) according to the manufacturer’s instructions. First-strand cDNA was synthesized by using SuperScript™ III First-Strand Synthesis System (Life Technologies Invitrogen, CA, USA) with oligo (dT)18 primer. The qPCR analysis was carried out by using DyNAmo ColorFlash SYBR Green qPCR Kit (Thermo Scientific^™^, USA) on a LightCycler-^®^ 480 Instrument II (Roche). The relative gene expression was normalized using *eef1a1l1* as an internal control. Oligonucleotide sequences for qPCR analysis are listed in Figure 6-source data 1.

### Immunostaining and imaging

For immunofluorescence, slides were washed twice with PBS and twice with ddH2O, and then incubated in blocking solution [1× PBS, 2% (vol/vol) sheep serum, 0.2% Triton X-100, 1% DMSO]. Slides were incubated in primary antibodies overnight at 4°C, followed by three PBST (0.1% Triton X-100 in 1× PBS) washes and incubation with secondary antibodies for 1.5 hr at 28°C. Slides were washed again with PBST, and stained with DAPI (Santa Cruz Biotechnology, Texas, USA) before mounting. Antibodies and reagents used in this study included anti-Mpx (Genetex, San Antonio, TX) at 1:500 and anti-mCherry (Abcam, Cambridge, UK) at 1:250. EdU staining was performed by using the Click-iT EdU Imaging Kit (Molecular Probes, Eugene, OR) following the manufacturer’s instructions. EdU (25 μg/fish) was IP injected 24 hr before extraction of the heart. TUNEL assay was performed by using *In Situ* Cell Death Detection Kit, TMR red (Sigma, St Louis, MO, USA) following the manufacturer’s instructions.

Imaging of whole-mount hearts and heart sections was performed by using Nikon SMZ25 and Zeiss LSM 800, respectively. Quantification of CM proliferation and CM density in zebrafish were performed in the 200 μm area directly adjacent to the injured area as previously described (19). Revascularization was examined by the endogenous fluorescence from live imaging. Quantification of re-vascularized density was measured from the whole injured area of the hearts and calculated by ImageJ. The student’s t-test was applied to assess all comparisons by Prism 9.

### Next-generation RNA sequencing analysis

For each time point and condition, zebrafish ventricles from 3 fish hearts were used as biological duplicates for the RNA-seq experiments. RNA extraction was performed as previously described with minor modifications (1). Briefly, RNA isolation was done by using the miRNeasy micro Kit (Qiagen). RNA quality analysis was done by using Qubit and Bioanalyzer at the NGS High Throughput genomics core, Academia Sinica. Sequencing was performed on the HiSeq Rapid (Illumina) setup, resulting in a yield of 15-20 M reads per library on a 2 x 150 bp paired-end setup at the NGS core facility. Raw reads were assessed after Quality control (QC), and output to filtered reads which were mapped to zebrafish Ensembl genome assembly using HISAT2 (20). The number of reads was aligned to Ensembl annotation using StringTie (21) for calculating gene expression as raw read counts. The raw counts of the mapped annotated genes were joined to a combined matrix and normalized using upper-quantile normalization to generate normalized Reads Per Kilobase of transcript, per Million reads mapped (RPKM) counts. Further analysis was done based on the normalized read values and respective log2 fold change to generate differentially expressed genes (DEGs) using NOIseq (22). Gene ontology (GO) enrichment analysis was performed in WebGestalt (23) for the dataset considering the upregulated and downregulated genes against control versus treatment conditions and respective uninjured samples. Over-representation analysis was done with id of mapped genes for multiple testing corrections using Benjamini and Hochberg FDR correction were conducted with a significance level of 0.05. Principle component analysis (PCA) was performed on Fragments Per Kilobase of transcript, per Million reads mapped (FPKM) normalized counts of all the datasets at respective time points to analyze the sample level progression through time under control versus treatment conditions. Pathway analysis was performed using Ingenuity Pathway Analysis (IPA) software (Qiagen) (24) following manufacturer protocols. Log2 ratio was input from the normalized read counts in zebrafish and defined as DEGs at log2FC above ±1.

### Single-cell RNA-sequencing (scRNAseq) and bioinformatic analysis

Heart dissociation was followed the same procedures in qPCR. For each scRNA-seq sample, 35-45 cryoinjured ventricles were collected from each experiment at respective time-points and conditions, pooled together and digested using LiberaseDH (Sigma, St Louis, MO, USA) at 28°C in a 24-well plate. Cell suspension was filtered through 100 μm, 70 μm and 40 μm cell strainers (SPL, Gyeonggi-do, Korea) and centrifuged at 200g for 5 min. Cell pellet was resuspended in 0.04% BSA/PBS, stained with DAPI and filtered through 35 μm Flowmi cell strainer (BD, New Jersey, USA). Cells were sorted by FACS (BD Facs Aria) and subjected to single-cell sequencing (scRNAseq) following cell counting with countess II automated cell counter (Invitrogen).

scRNAseq library was generated using single cell 3’ Reagent kits in chromium platform (10x Genomics). Cell ranger software suite (10x Genomics) was utilized for processing and de-multiplexing raw sequencing data (25). The raw base files were converted to the fastq format and the subsequent sequences were mapped to zebrafish Ensembl genome assembly for processing. Downstream analysis of the gene count matrix generated by CellRanger (10x Genomics) was performed in R version of Seurat 3.1 (26, 27). The raw gene count matrix was loaded into Seurat and a Seurat object was generated by filtering cells that expressed more than 400 nUMI counts and over 200 genes. Additional filters for extra quality control was applied by filtering cells with log10genes perUMI > 0.8 and cells with mitochondrial gene content ratio lower than 0.23. This resulted in 9437 cells for the uninjured dataset, 16657 cells for the PBS1d dataset, 9997 cells for PBS3d dataset, 11950 cells for PBS7d dataset, 9912 cells for CL1d dataset, 12621 cells for the CL3d dataset and 11236 cells for the CL7d dataset. Reads were normalized by the “NormalizeData” function that normalizes gene expression levels for each cell by the total expression. The top 3000 highly variable genes were used as variable features for downstream analysis. Prior to dimensionality reduction, a linear transformation was performed on the normalized data. Cells were scored by “Cellcyclescoring” function and Unwanted cell-cell variation was eliminated by “regress out’ using mitochondrial gene expression during scaling. The datasets were integrated into a single Seurat object by using the canonical correlation analysis (CCA) with a “SCT” normalization method. Dimensionality reduction was performed on the integrated dataset using PCA. The top 40 principal components were identified based on PCElbowPlot. UMAP dimensional reduction by “RunUMAP” function was used to visualize the cell clusters across conditions (28) (Figure 2-source data 1). 19 cell clusters were generated by “FindClusters” function using 40 PCs and a resolution of 0.4. Following clustering, DEGs in each of the clusters were determined using “FindMarkers” and “FindAllMarkers” assessed by the statistical MAST framework (29). Top DEGs of each cluster were then filtered based on being detected in ≥25% of cells compared to other cells within the dataset, with a Bonferroni adjusted p-value <0.05. Clusters were assigned with specific cell identities based on the lists of DEGs and ordered by average log fold change and p-value. Visualization of specific gene expression patterns across groups on UMAP and heatmaps (top enriched genes) was performed using functions within the Seurat package. Gene enrichment analysis were determined using the above described RNA-seq pipeline. In brief, the DEGs across cell types were used in WebGestalt to generate the cluster-specific biological processes of GO and KEGG pathways (30).

### Pseudobulk analysis of scRNAseq

We generated reference matrixes of the unsupervised and un-normalized mean counts computed for each gene across all individual cells from each cell type using the Seurat and the SingleCellExperiment package (31). The raw matrices were split by cell type and the associated conditions of zebrafish (PBS-treated vs CL-treated). Normalization was performed on the sparse gene matrix using total read counts and the total cell numbers for each cell type. The normalized count matrix was applied for differential expression analysis by using NOISeq (22). Furthermore, average expression of genes across all the cell types detected by the scRNA-seq data from each time point was calculated using Seurat function “AverageExpression”, which was used as a reference dataset for query. The hierarchical heatmap was generated by Morpheus (https://software.broadinstitute.org/morpheus).

### Ligand-Receptor Interaction Network Analysis

The zebrafish ligand-receptor pairs were derived from Human ligand-receptor pair database as described previously (32). We mapped the ligand-receptor pair orthologous to the human database and generated the fish database for use in interaction network. We kept the ligand-receptor pairs that showed expression in upper-quantile normalization as the cutoff to define the expressed data. Next, the macrophages and neutrophils that expressed the genes of ligands and receptors were determined based on the counts from pseudobulk analysis (which was described in the section of Pseudobulk analysis of scRNAseq) with an upper-quantile expression at the given time point. The total number of the potential ligand-receptor interaction events between each cluster of macrophages and neutrophils were visualized using Cytoscape (33).

## Results

### Delayed macrophage recruitment results in prolonged inflammation and disrupted energy metabolism during cardiac repair

In zebrafish, macrophage pre-depletion at 1-day prior to cardiac injury (−1d_CL) delayed macrophage recruitment and compromised heart regeneration, even though the overall macrophage numbers were restored within the first week (1). To investigate the global transcriptomic changes under these regenerative (PBS-treated, normal macrophage recruitment) and non-regenerative (CL-treated, delayed macrophage recruitment) conditions, we isolated zebrafish hearts at 7 and 21 day-post cryoinjury (dpci), corresponding to the time when cardiomyocytes proliferate and replace the scar tissue during heart regeneration, and subjected the hearts to bulk RNAseq analyses (Figure 1A). We first plotted our data from 7 and 21 dpci with published data points from 6 hours post cryoinjury (hpci) to 5 dpci in a Principal Component Analysis (PCA, Figure 1B). Transcriptomes of the PBS-treated samples at 7 and 21 dpci nicely fit into the trajectory between 5 dpci and untouched hearts, suggesting that transcriptomic changes in control hearts coincide with the cardiac repair toward the naïve state (purple dots and trail, Figure 1B). However, the transcriptome of non-regenerative CL-treated hearts at 21 dpci (CL21d) was in proximity to that of PBS-control hearts at 7 dpci (PBS7d), suggesting a delayed transcriptomic response in the non-regenerative CL-treated hearts (green dots, Figure 1B).

**Figure 1.**
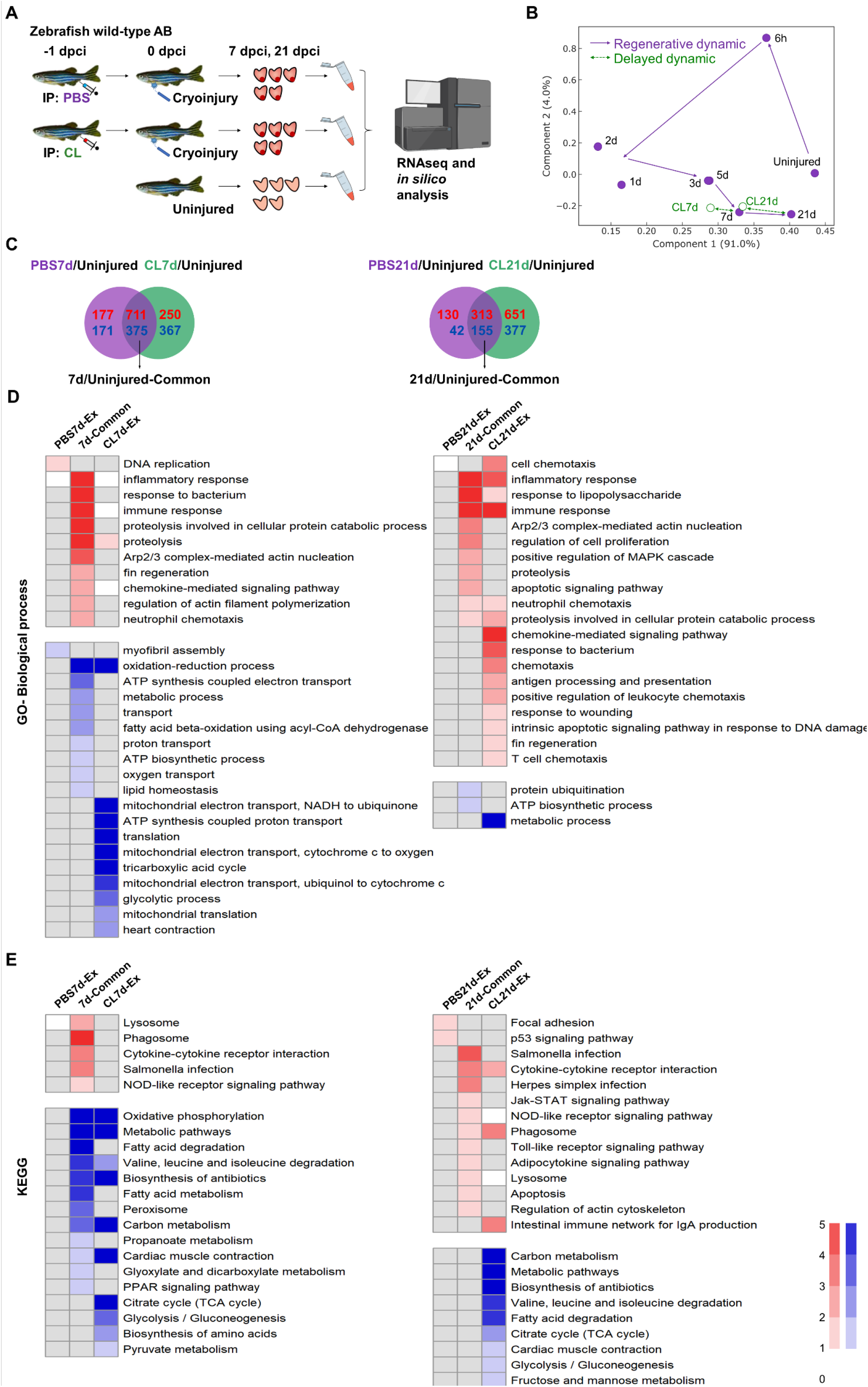
Transcriptional profiling of infarcted hearts under regenerative and non-regenerative conditions. (A) Experimental design. Zebrafish AB_wildtype was IP-injected with PBS or Clodronate liposomes (CL) one-day before cardiac cryoinjury. Injured hearts were collected at 7 and 21 day post cryoinjury (dpci), respectively. Uninjured hearts were collected as the control of the baseline. Total RNA was extracted and analyzed by RNA sequencing (RNAseq). (B) Principal Component Analysis (PCA) of gene expression in hearts at different time points. The PCA was performed on the FPKM normalized datasets of healthy hearts and injured hearts at 7 and 21 dpci after PBS or CL treatments (PBS7d, PBS21d, CL7d and CL21d). The datasets of 0h, 6h, 1d, 2d, 3d, and 5 dpci from the previous study were also included^1^. Regeneration and delayed dynamics were indicated by purple and green lines, respectively. FPKM, Fragments Per Kilobase of transcript per Million mapped reads. (C) Venn diagrams showed the comparatively differentially expressed genes (DEGs). Up-regulated and down-regulated gene numbers were highlighted in red and blue, respectively. Exclusive up-regulated and down-regulated genes at 7 dpci (left panel) and 21 dpci (right panel) in PBS-treated hearts (regenerative condition) were shown in purple, while the counterparts in CL-treated hearts (non-regenerative condition) were shown in green. (D, E) Gene Ontology (GO) and Kyoto Encyclopedia of Genes and Genomes (KEGG) pathway analyses of differentially expressed genes (DEGs). The enrichment of DEGs at 7 and 21 dpci in PBS-treated or CL-treated hearts was analyzed by GO and KEGG pathway. Significant values for the enriched pathways of up- and down-regulated genes were shown in red and blue, respectively. Gene expression in healthy hearts was used as the baseline control. Statistics were performed using a hypergeometric test (Fisher’s exact test; P<0.05) and the scale bar showed significant values as −log10 (P_adj_).

We next defined the differentially expressed genes (DEGs) by comparing gene expression levels in PBS-treated or CL-treated hearts against their basal expression levels in untouched hearts, and then divided them into PBS-specific, CL-specific, and common DEGs at 7 and 21 dpci using Venn diagrams (Figure 1C). At 7 dpci, the number of upregulated DEGs was comparable between PBS and CL-treated hearts, while more downregulated DEGs were identified in CL-exclusive conditions. The number of DEGs decreased drastically from 7 to 21 dpci in PBS-control hearts, while the number remained high in CL-treated hearts, suggesting a retained and/or aberrant response corresponding to the PCA analysis (Figure 1C). Common and condition-specific DEGs were then subjected to gene ontology (GO) and Kyoto Encyclopedia of Genes and Genomes (KEGG) pathway analyses (Figure 1D and 1E). At 7 dpci, up-regulated DEGs in both PBS- and CL-treated hearts were associated to “immune/inflammatory response”, “chemokine-mediated signaling”, “proteolysis”, “neutrophil chemotaxis”, “phagocytosis”, “cytokine-cytokine receptor interaction”, and “NOD-like receptor signaling pathway” (7d-Common, Figure 1D and 1E). These processes correspond nicely with known repairing processes of immune cell infiltration and debris clearance in the first week of cardiac repair. At 21 dpci, CL-exclusive DEGs were still enriched in “immune/inflammatory response”, particularly in “cytokine/chemokine and chemotaxis-related pathways”, suggesting a prolonged inflammation and cell infiltration of the injured tissue (CL21d-Ex, Figure 1D and 1E). On the other hand, downregulated DEGs in both PBS and CL-treated hearts were enriched in “oxidative phosphorylation” and metabolism at 7 dpci and restored at 21 dpci, corresponding to the metabolic shift of energy production from fatty acid oxidation to glycolysis during cardiac repair and hypoxia (Figure 1E) (34). Interestingly, downregulated CL-exclusive DEGs were enriched in glycolytic pathways including “TCA cycle”, “Carbon metabolism”, and “Glycolysis/Gluconeogenesis” at both 7 and 21 dpci (CL21d-Ex, Figure 1D and 1E). Taken together, these results indicate that CL treatment prolonged the inflammatory phase of the repairing process, which normally occurs only in the first week post injury and interfered with energy metabolism during cardiac repair.

To highlight the DEGs that behave differently under regenerative PBS vs. non-regenerative CL conditions, we performed hierarchical clustering of all DEGs (Figure 1-figure supplement 1 and Figure 1-figure supplement 1-source data 1). These DEGs were grouped into 22 clusters with similar expression dynamics corresponding to time points and treatments. Among regeneration-associated clusters, genes within the Cluster 2 (C2) were up-regulated only in PBS-hearts at 7 dpci, including genes in “response to oxidative stress” (*apodb, apoda.1, anxa1a*, and *hmox1a*) and other unclassified genes (*serpine1, havcr2, tnfaip6*, and *stmn1a*). These genes may contribute to heart regeneration by ROS regulation and inflammatory modulation. For example, *Serpine1* re-activates in wound endocardium and is involved in endocardial maturation and proliferation, fibrinogenesis, and CM proliferation during heart regeneration (35, 36). *havcr2* encodes an inhibitory receptor TIM-3 found in a variety of immune cells and regulates both innate and adaptive immune responses, including apoptotic body clearance, nucleic acid–mediated innate immune responses, and T cell survival (37). *tnfaip6* also encodes an important immunoregulatory factor TNFα induced protein 6 to activate macrophage transition from a pro-inflammatory to an anti-inflammatory phenotype (38, 39). The DEGs in C9-C12 were up-regulated only in PBS-control hearts at 21 dpci, containing several ribosomal subunit genes and suggesting active production of building blocks for replenishing lost tissues (Figure 1-figure supplement 1). Furthermore, DEGs in C19 and C20 function in various metabolic processes, including “Carbon metabolism”, “Fatty acid degradation” and “Glycolysis/Gluconeogenesis”, which were active in uninjured hearts, downregulated at 7 dpci, and reactivated only in PBS-treated but not CL-treated hearts at 21 dpci (Figure 1-figure supplement 1). CMs undergo metabolic changes from anaerobic to oxidative metabolism during differentiation and maturation to meet the high energy demands from constant beating (40). Upon myocardial infarction and heart failure, adult CMs switch back to using glucose (glycolysis) instead of fatty acid (oxidative phosphorylation) as the main substrate for energy (34, 41). Interestingly, this metabolic switch was also observed during zebrafish cardiac repair and regeneration, when pyruvate metabolism and glycolysis are beneficial for CM dedifferentiation and proliferation (42). As most of these processes take place in mitochondria, cardiac resident macrophages maintain mitochondrial homeostasis of healthy myocardium by eliminating dysfunctional mitochondria from CMs (43), while mitochondrial dysfunction leads to impaired revascularization and fibrosis formation in mammals (44–46). Nevertheless, direct involvement of macrophages in regulating metabolic shift upon cardiac injury has not been yet demonstrated. Further studies are required to test whether mis-regulated metabolism is associated with delayed macrophage recruitment in CL-treated hearts.

**Figure 1-figure supplement 1.**
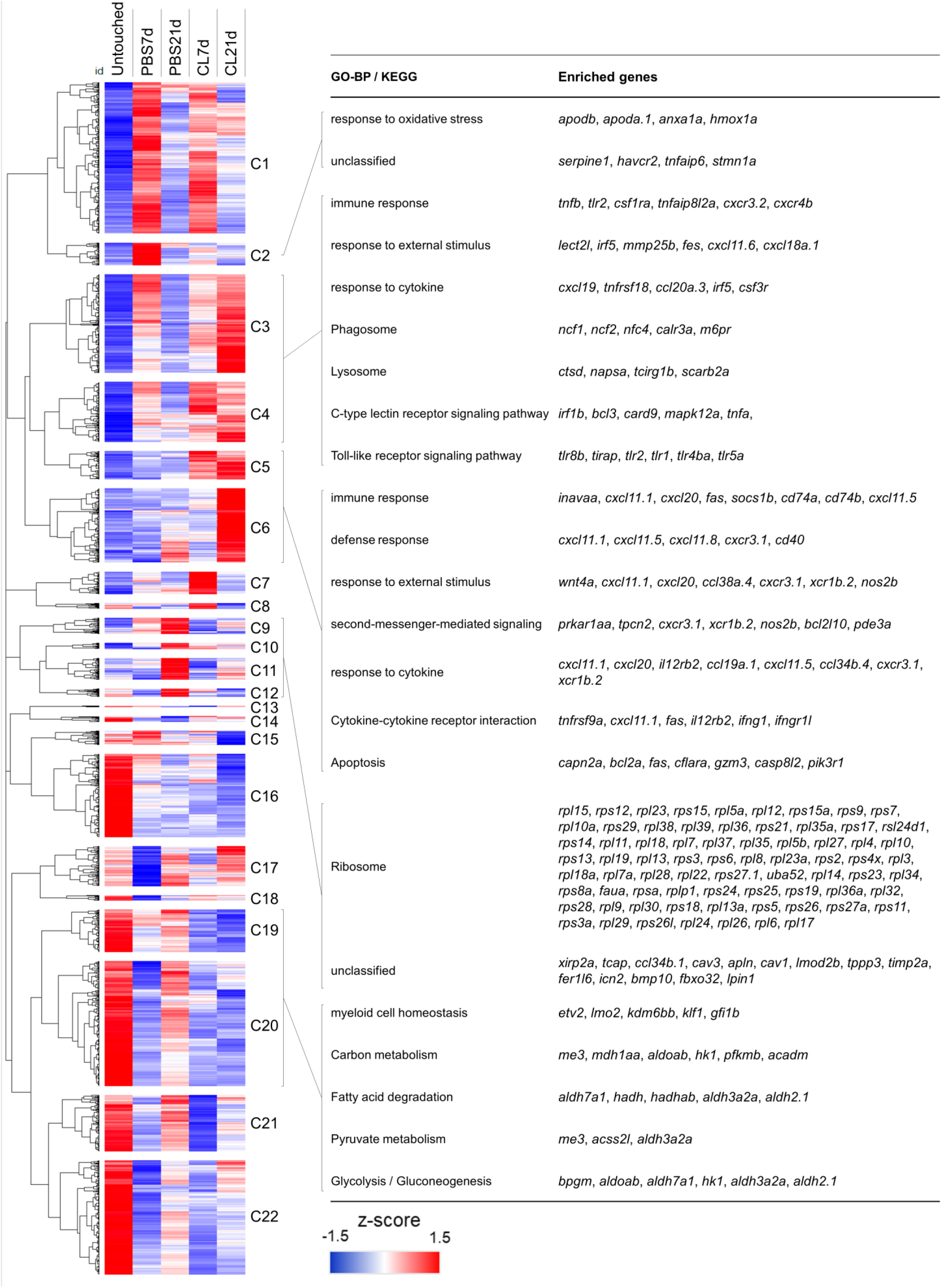
Hierarchical clustering heatmap of the DEGs under regenerative and non-regenerative conditions in zebrafish. Heatmap of all DEGs selected by NOIseq (q>0.99) and arranged by hierarchical clustering from cluster 1 (C1) to cluster 22 (C22) (left panel). The value was a z-score from 1.5 as red to −1.5 as blue. BP of GO and KEGG pathways of the DEGs were analyzed by using WebGestalt (right panel). Cluster-enriched genes involved in their predicted biological processes and pathways were listed. The threshold of enriched categories was FDR<0.05. DEGs, differentially expressed genes; GO, Gene Ontology; BP, biological process; KEGG, Kyoto Encyclopedia of Genes and Genomes; FDR, False discovery rate.

Among the clusters associated with non-regenerative CL condition, C3 and C4 contain genes generally activated at 7 dpci in both PBS and CL-hearts, but their activation was prolonged and even intensified in CL-treated hearts at 21 dpci (Figure 1-figure supplement 1). The biological functions of these DEGs are mainly related to immune response, including “response to external stimulus and cytokine”, “C-type lectin receptor signaling pathway” and “Toll-like receptor signaling pathway”. (Figure 1-figure supplement 1). Prolonged inflammation may prevent tissue repair and regeneration (47). Inflammation also induces continuous neutrophil migration and infiltration and prevents their clearance by apoptosis (48). Correspondingly, up-regulated DEGs involved in “immune response”, “Cytokine-cytokine receptor interaction”, and “Apoptosis” appeared at 21 dpci in CL-treated hearts (C5 and C6). Prolonged activation of apoptosis indicates that cardiac cells (including CMs) continuously undergo programmed cell death potentially suffering from ROS stress (ferroptosis) and inflammatory microenvironment (pyroptosis) (49, 50). Continuous cell death subsequently triggers the prolonged activation of C-type lectin receptor singling pathway in active phagocytes (51, 52). As a result, proteolysis of engulfed debris and cell adhesion molecules were activated under the same conditions (Figure 1-figure supplement 1). Altogether, the results of hierarchical clustering nicely reflect the GO analysis and suggest that CL treatment disrupts inflammatory resolution and energy metabolism during cardiac regeneration. The full list of DEGs in each cluster was summarized in Figure 1-figure supplement 1-source data 1.

To identify the canonical pathways and potential upstream regulators associated with aberrant regeneration, these DEGs were further analyzed by Ingenuity Pathway Analysis (IPA) (Figure 1-figure supplement 2). CL treatment led to continuous activation of “Leukocyte Extravasation Signaling” and “Production of NO and ROS in Macrophages” pathways (Figure 1-figure supplement 2A). Correspondingly, downstream genes of inflammatory cytokines IFNG, TNF, and IL6 were continuously activated in CL-treated hearts until 21 dpci, while they were largely down-regulated in PBS-control hearts at 21 dpci (Figure 1-figure supplement 2B). Among genes of the “Leukocyte Extravasation Signaling” pathway, we found several integrin genes usually expressed on the leukocyte plasma membrane, including *itga4, itgal* and *itgb2*, which are involved in leukocyte enrolling on endothelial cells and transmigration (53, 54). In addition, we found several matrix metalloproteinases (MMPs) including *mmp9, mmp13*, and *mmp25*, which might be involved in ECM remodeling and leukocyte recruitment during inflammation (Figure 1-figure supplement 2C) (55–57). These results support our previous observation on continuous neutrophil infiltration and retention in the CL-treated hearts even until 30 dpci (1). Lastly, among genes of the “NO and ROS production in Macrophages” pathway, we found continuous activation of DAMP/PAMP receptor *tlr2*, neutrophil cytosolic factors (*ncf1, ncf2*, and *ncf4*), myeloid cell-lineage committed gene *spi1* and its downstream target *ptpn6*, in addition to the macrophage differentiation marker *irf8* in CL-treated hearts at 21 dpci (Figure 1-figure supplement 2D). ROS play dual roles in tissue repair and are largely produced by macrophages (58). On one hand, ROS attract leukocytes recruitment to the wound (59) and promote CM proliferation post cardiac injury (60). On the other hand, uncontrolled production of NO and ROS may be detrimental, leading to mitochondrial damage (61), blocking macrophage differentiation from monocytes to anti-inflammatory phenotype (62–64), and extending immune response (65, 66). Collectively, these results indicate that macrophages may play roles in regulating ROS homeostasis, immune cell dynamics, and inflammation resolution, as these processes were mis-regulated under CL treatments.

**Figure 1-figure supplement 2.**
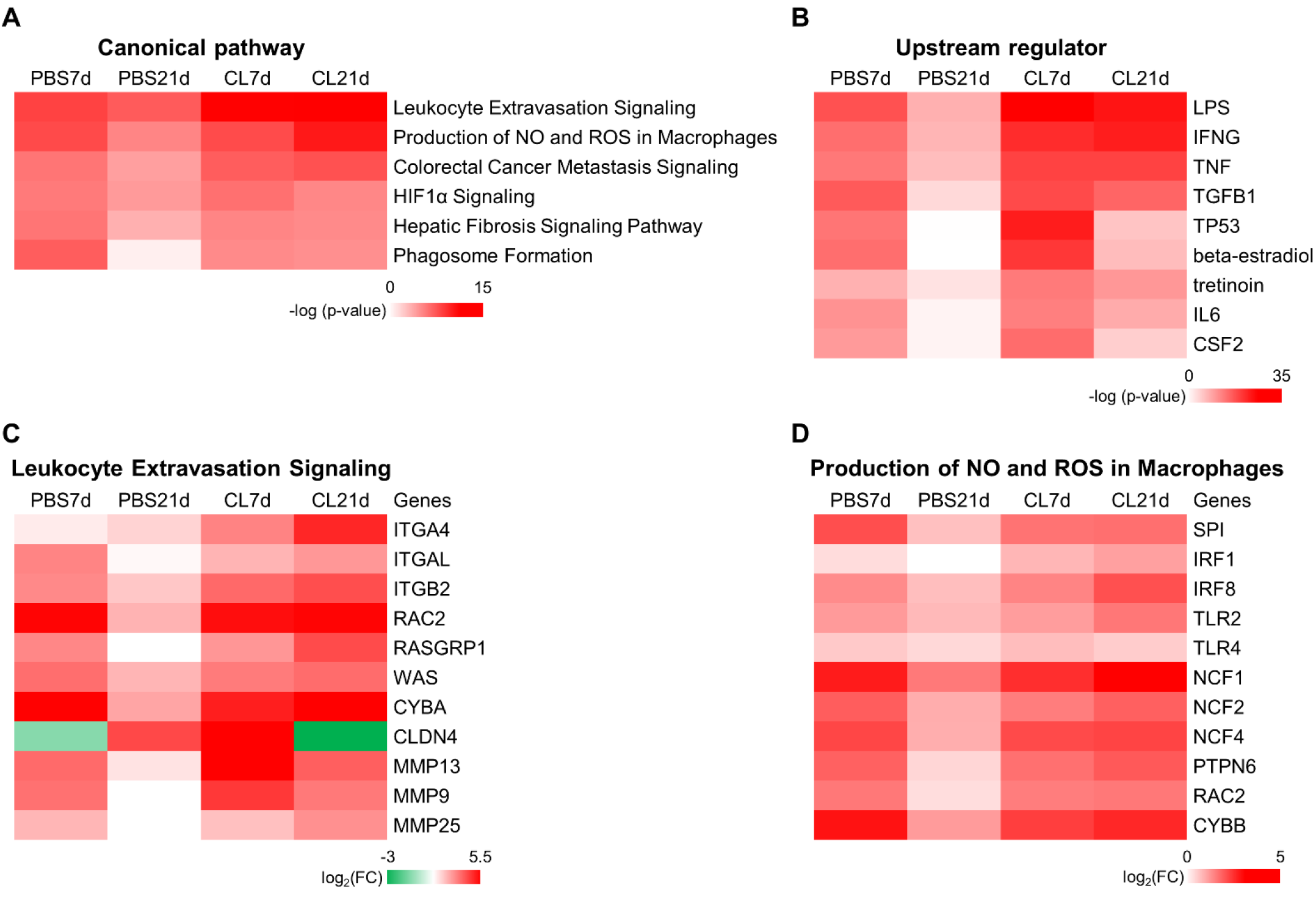
Identification of canonical pathways and upstream regulators under regenerative and non-regenerative conditions in zebrafish. (A and B) Top canonical pathways (A) and upstream regulators (B) in PBS-treated or CL-treated hearts were predicted by Ingenuity Pathway Analysis (IPA). The value was shown as −log(p-value) and the color reflected the degree of enrichment. (C, D) The heatmaps of top DEGs involved in the canonical pathways of Leukocyte Extravasation Signaling (C) and Production of NO and ROS in Macrophages (D). The value was Log2 fold change indicated by color at bottom. DEGs, differentially expressed genes.

### Single-cell analyses reveal the heterogeneous landscape of inflammatory cells in the infarcted hearts under regenerative and non-regenerative conditions

Since macrophage properties may be altered upon CL-treatment and thus fail to support heart regeneration by resolving neutrophil infiltration and inflammation, we next analyzed and compared the potential identity and function of these inflammatory cells by single-cell transcriptomic profiling (Figure 2A). Double transgenic zebrafish *Tg(mpx:EGFP;mpeg1:mCherry)* were IP injected with CL or PBS at 1 day before cryoinjury, and the EGFP expressing neutrophils and mCherry expressing macrophages were isolated by fluorescence-activated cell sorting (FACS) from the uninjured hearts, as well as regenerative (PBS-control) and non-regenerative (CL-treated) hearts at 1, 3 and 7 dpci (Figure 2A and Figure 2-figure supplement 1A) (13, 67). In the uninjured hearts, we found a substantial number of mCherry^+^ and mCherry^+^EGFP^+^ resident cells (mostly macrophages, ~0.6% of total cardiac cells) and very few EGFP^+^ neutrophils (Figure 2-figure supplement 1A). Among them, mCherry^+^EGFP^+^ cells show higher complexity and larger size (FCS-A and SSC-A) than mCherry^+^ cells, corresponding to the macrophages and the progenitor/lymphocyte properties previously described (Uninjured sample in Figure 2-figure supplement 1A) (68). After injury, both macrophages and neutrophils increased rapidly, and divergent cell composition and numbers were observed in PBS vs. CL-treated hearts over time. While macrophage numbers kept increasing after cardiac injury until 7 dpci, neutrophils peaked at 3 dpci and gradually decreased between 3 to 7 dpci in PBS-control hearts, corresponding to the inflammatory resolution phase previously described (9). In CL-treated hearts, macrophages progressively increased, similar to control hearts, but neutrophil numbers became much higher than controls at both 3 and 7 dpci (Figure 2-figure supplement 1A). At 7 dpci, a similar percentage of macrophages were sorted under both conditions, while a higher percentage of neutrophils were sorted in CL conditions (Figure 2-figure supplement 1B). These results coincide with published findings and imply that macrophage numbers had recovered, but neutrophil resolution was delayed in CL-treated hearts compared to PBS-control hearts (1). Interestingly, those EGFP^+^/mCherry^+^ macrophages resided nearby the epicardial layer of the uninjured/naïve hearts and proliferated to maintain their population, similar to murine resident macrophages (Figure 2-figure supplement 1C and D) (69). Upon cardiac injury, those resident macrophages were preferentially enriched in regenerative hearts at 1 dpci to 3 dpci (Figure 2-figure supplement 1A) and populated the infarct area at 7 dpci (Figure 2-figure supplement 1E).

**Figure 2.**
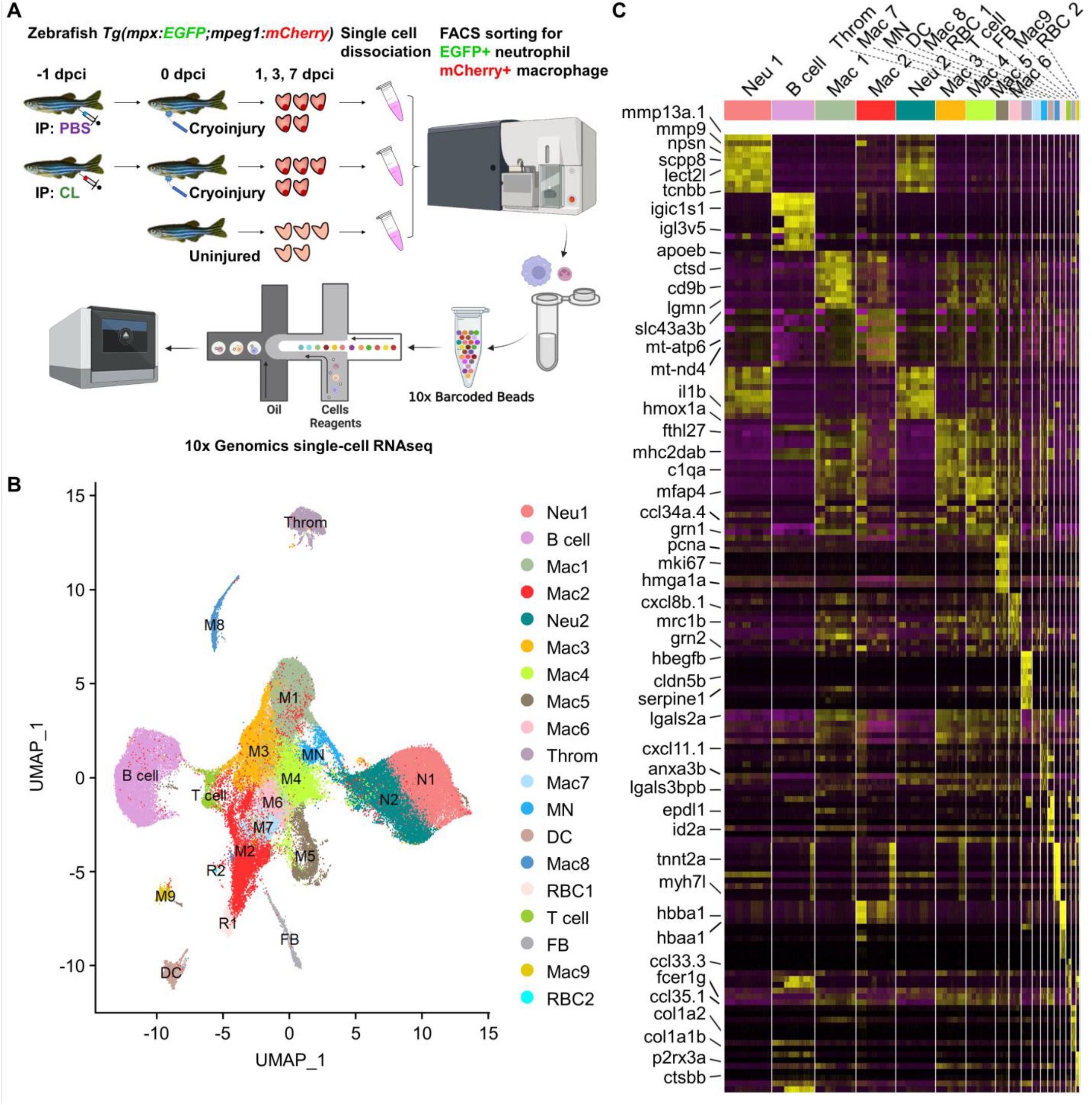
Temporal single-cell analyses revealed heterogenous macrophages and neutrophils in the infarcted hearts. (A) Experimental design. Double transgenic *Tg(mpx:EGFP;mpeg1:mCherry)* zebrafish expressing EGFP in neutrophils and mCherry in macrophages were IP-injected with PBS (regenerative condition) or CL (non-regenerative condition) one day before cryoinjury (−1 dpci, −1d_CL). Injured hearts were collected and dissociated into single cells at 1, 3 and 7 dpci. Untreated and uninjured hearts were also collected and dissociated. Single cells of each time point were then subjected to a fluorescence-activated cell sorter (FACS) for isolating the mCherry^+^ and EGFP^+^ cells. RNA was purified from these cells and barcoded followed by single-cell RNA sequencing (scRNAseq). (B) Uniform Manifold Approximation and Projection (UMAP) of the isolated cells. The isolated cells consisted of 9 macrophage clusters, 2 neutrophil clusters, 1 hybrid cluster (MN), and other minor populations including B cell, thrombocyte, dendritic cell (DC), T cell, fibroblast (FB), and 2 red blood cell (RBC) clusters. (C) Heatmap of top 10 DEGs in 19 clusters of infarcted hearts. yellow highlights the enriched genes. Cluster-enriched/specific genes are listed on the left.

**Figure 2-figure supplement 1.**
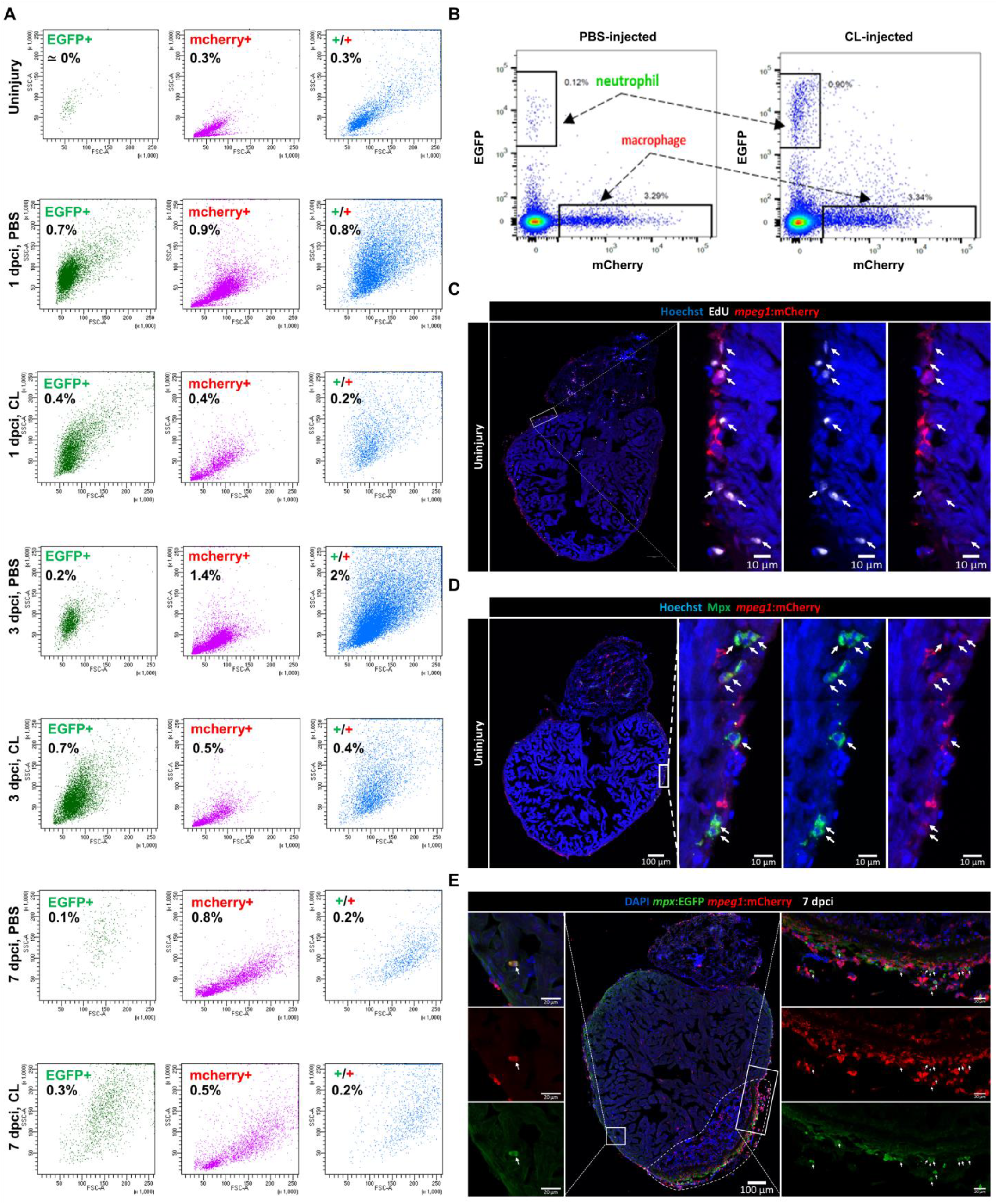
Dynamics of macrophages and neutrophils in zebrafish heart prior and post injury. (A) Macrophages and neutrophils derived from uninjured and infarcted hearts under regenerative (PBS) or non-regenerative (CL) conditions were analyzed by FACS. Cell granularity and size were shown by side scatter (SSC)/ forward scatter (FSC) gating. Neutrophils (EGFP^+^ cells) were shown in green, macrophages (mCherry^+^ cells) were shown in pink, and double-positive cells were shown in blue. Cell proportion of total input was indicated in each panel. (B) Representative plots of the EGFP and mCherry gating of 7 dpci PBS and CL sorted macrophages and neutrophils. (C) Examination of steady-state macrophage proliferation locally in hearts by Edu staining. White arrows highlighted the proliferating (EdU^+^mCherry^+^ double-positive) cells. (D) Validation of Mpx^+^ macrophages in uninjured *Tg(mpeg1:mCherry)* heart cryosection by using anti-Mpx antibody. White arrows highlighted the Mpx^+^mCherry^+^ double-positive cells. (E) Validation of mpx^+^ macrophages on 7 dpci *Tg(mpx:EGFP;mpeg1:mCherry)* heart cryosection. White arrows highlighted the mCherry^+^EGFP^+^ double-positive cells.

All these inflammatory cells were then subjected to droplet-based high-throughput single-cell RNA sequencing (scRNAseq). To visualize the dataset, sequencing reads were mapped to the zebrafish genome, assigned to each cell, and then processed by the Uniform Manifold Approximation and Projection (UMAP) for dimension reduction and unbiased clustering using Seurat R package (Figure 2B) (27). Quality control was conducted according to the pipeline in Seurat based on the read numbers per cell (nUMI), the number of genes detected per cell (nGene), and mitochondrial counts ratio to keep the cells of high quality for downstream analyses (Figure 2-figure supplement 2A-D). This resulted in 9,437 cells for uninjured dataset, 16,657 cells for PBS1d dataset, 9,997 cells for PBS3d dataset, 11,950 cells for PBS7d dataset, 9,912 cells for CL1d dataset, 12,621 cells for CL3d dataset and 11,236 cells for CL7d dataset (Figure 2-figure supplement 2E). After clustering, we identified 19 distinct clusters for these inflammatory cells and found that all of them expressed myeloid lineage marker genes *spi1b* and *coro1a* (Figure 2B and Figure 2-figure supplement 3A). Among them, 9 macrophage clusters (Cluster Macs), 2 neutrophil clusters (Cluster Neus), and one hybrid cluster (Cluster MN) were identified, based on the expression of reporter genes *mpeg1* and *mpx*, as well as other mononuclear phagocyte markers *csf1r, ccr2, cxcr1, irf8, lyz, mfap4*, and *kita* (Figure 2B and Figure 2-figure supplement 3A). We also identified small populations of B cells, T cells, dendritic cells, thrombocytes, red blood cells, and fibroblasts based on respective marker genes shown in UMAP and heatmap (Figure 2B and Figure 2-figure supplement 3B). Correspondingly, a subpopulation of B cells has been previously shown to express *mpeg1* and observed in *mpeg1:mCherry* fish (70). Minor *mpeg1^-^/mpx^-^* clusters might come from contamination during FACS, even though stringent gating strategies were applied, and will not be further analyzed in this study (71, 72).

Besides common lineage markers, macrophages showed cluster-enriched/specific gene expression (Figure 2C). As the biggest subpopulation, Mac 1 preferentially expressed *apoeb*, *ctsd*, *lgmn*, and *cd9b*, genes associated with lysosomal activity and phagocytosis. Mac 2 uniquely expressed hemoglobin-encoding genes such as *hbba1* and *hbaa1* that might have a role in protecting from nitric oxide (NO) or ROS (73). Mac 3 was enriched in *mhc2dab, c1qa* and *mfap4* that are associated with immune process, including antigen presentation and complement activation. In addition, Mac 3 expressed high levels *timp4.3* which might contribute to cardiac repair by modulating MMPs activities and ECM compositions (74–76). Mac 4 preferentially expressed *grn1, cxcl8a* and *hbegfb* that are associated with regeneration and growth as well as neutrophil recruitment (77, 78). Mac 5 was enriched in proliferating markers *mki67* and *pcna*, which is similar to a common macrophage subset identified and described in mammalian cardiac injury (72). Mac 6 expressed high levels of *cxcl8b.1, mrc1b* and *grn2*. Mac MN was enriched in *cxcl11.1*, which is a specific macrophage chemoattractant through the cxcl11.1-cxcr3.2 axis (79), as well as *lgals3bpb*, a gene enriched in microglia and predicted to have scavenger receptor activity (80). Mac 8 population was enriched in *myh7l* and *tnnt2a*, which are typical CM markers, and this result was in line with a previous study in mouse showing that cardiomyocyte signature is specifically expressed in murine cardiac CX_3_CR_1_^+^ resident macrophages (81). Mac 9 expressed *kita, tcf12*, and *csf1ra*, resembling the myeloid progenitors identified in mouse and zebrafish embryonic stage (82–84). Unlike heterogeneous macrophages, neutrophils were classified into just two populations Neu 1 and Neu 2, which commonly express *mmp9, mmp13a.1, npsn, scpp8, lect2l, tcnbb* and *il1b* genes (Figure 2C). *Mmp9*, as well as other granular protein and integrin genes-*mmp25, itgb2* and *gm2a*, were enriched in both clusters, suggesting they were both mature neutrophils (Figure 2-figure supplement 3C) (85). In addition, interferon-stimulated genes such as *rsad2, isg15, ifit8* and *mxa* were enriched in the Neu 2 and the *mpeg1*^+^ macrophages MN (Figure 2-figure supplement 3D). The full list of cluster-enriched genes were summarized in Figure 2-source data 2. These results revealed the heterogeneous landscape of inflammatory cell subpopulations in the zebrafish injured hearts, and further investigations are required to identify specific markers and examine the potential function of each cell cluster.

**Figure 2-figure supplement 2.**
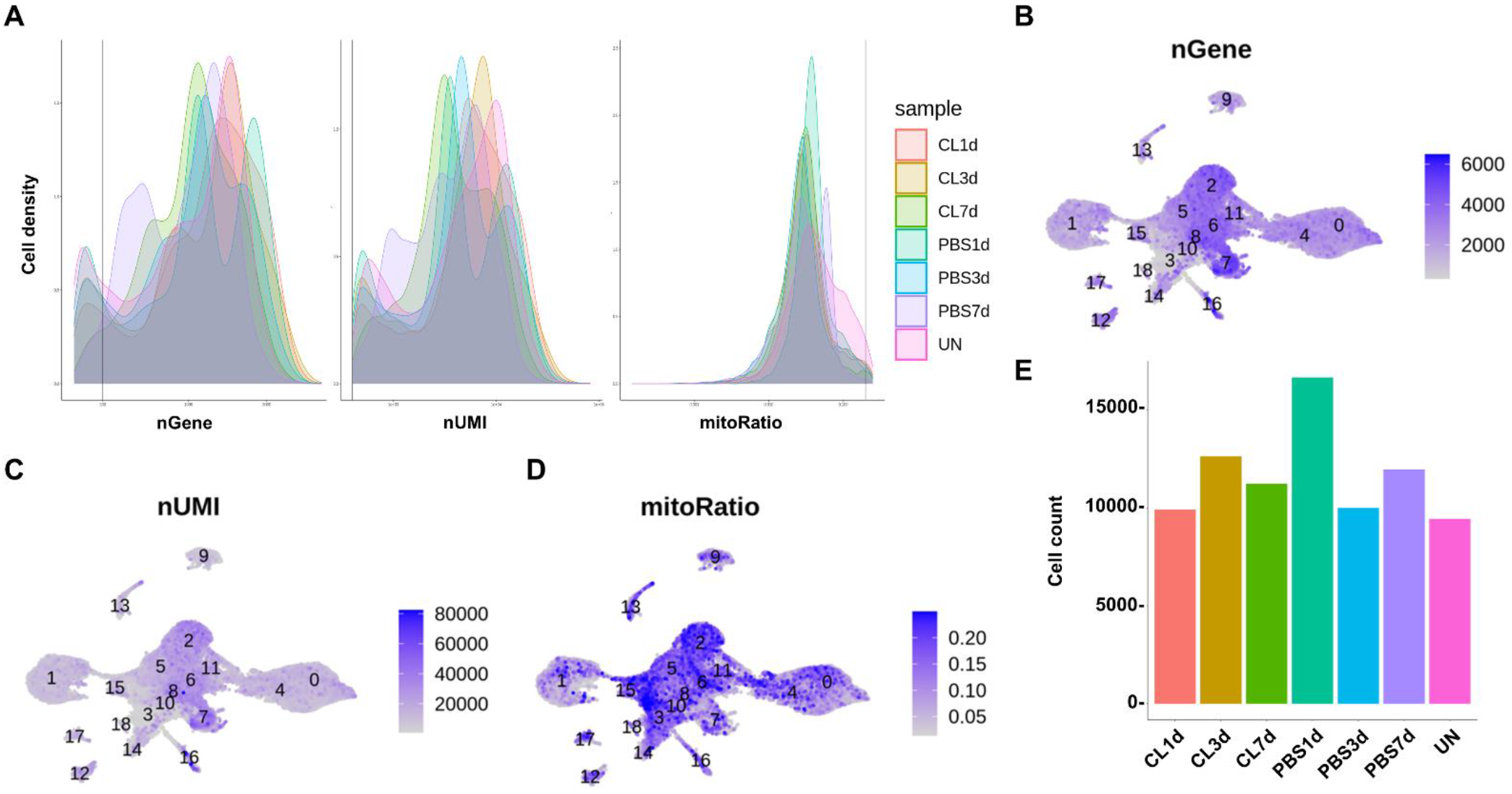
Quality controls for all the scRNAseq datasets by Seurat. (A) Respective plots of nGene, nUMI and mitoRatio versus cell density for each time point and condition. (B-D) UMAP plots colored by the counts of nGene (B), nUMI (C), and the percentage of mitoRatio (D). (E) Filtered cell counts after quality control for each time point and condition.

**Figure 2-figure supplement 3.**
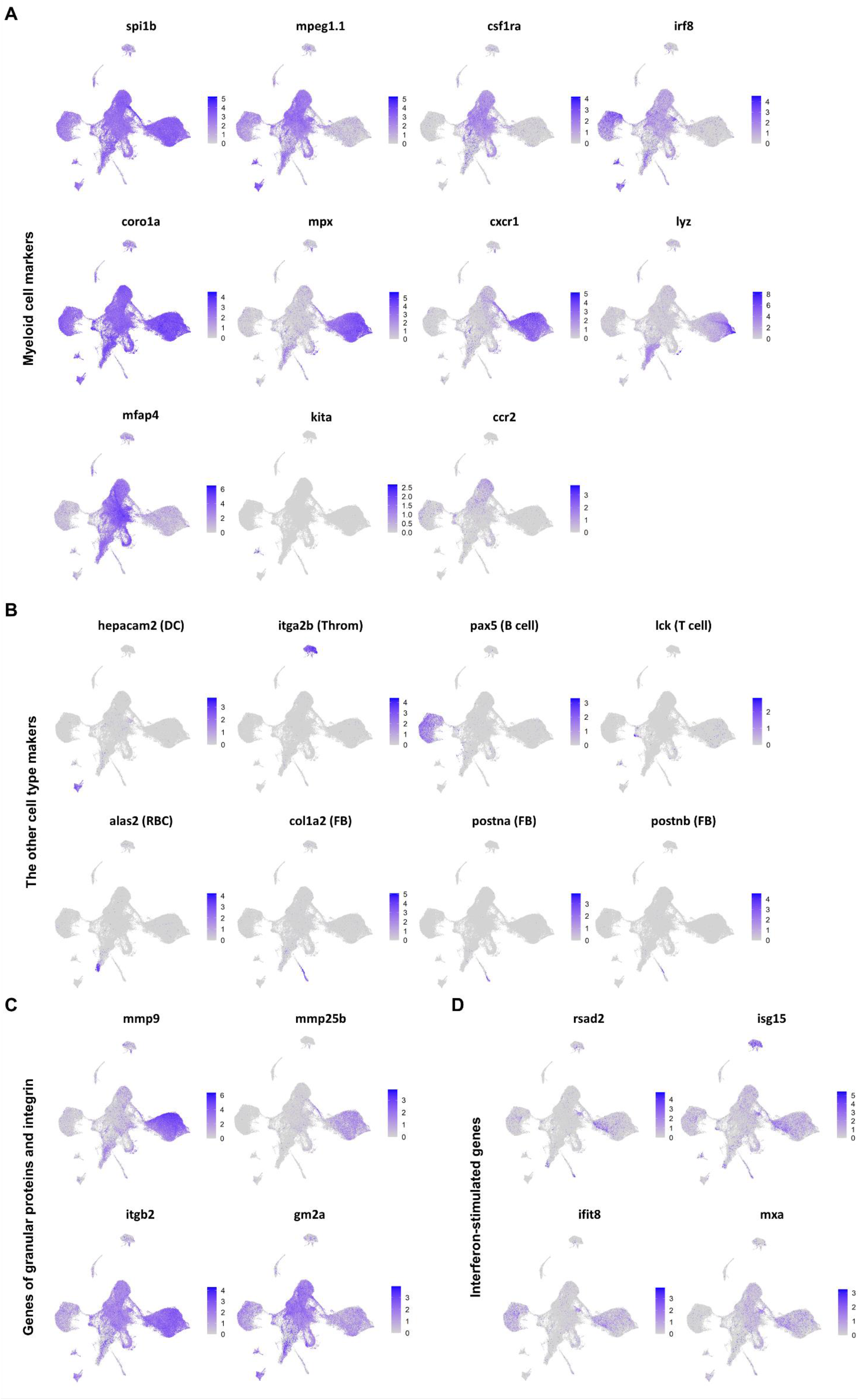
Expression of selected genes visualized on UMAP plots. (A) UMAP plots of macrophages and neutrophils marker genes. (B) UMAP plots of marker genes of dendritic cells (DC; *hepacam2*), thrombocytes (Throm; *itga2b*), B cells (*pax5*), T cells (*lck*), red blood cells (RBC; *alas2*) and fibroblasts (FB; *colla2, postna* and *postnb*). (C) UMAP plots of genes encoded granular proteins and integrin in mature neutrophils. (D) UMAP plots of interferon-stimulated genes (ISGs).

### Dynamic cell proportion analyses of inflammatory cell clusters associate specific resident macrophage subsets with heart regeneration

To dissect the dynamic changes of these inflammatory cell clusters under regenerative vs. non-regenerative conditions, we generated split UMAPs for each time point and condition (Figure 3A). In uninjured heart/naïve state, Mac 2 and 3 represented the major resident macrophage clusters followed by Mac 1, 4 and 8 (Figure 3B). The proliferating macrophage cluster Mac 5 was also observed in the uninjured hearts, corresponding to those residing nearby the epicardial tissue, suggesting that some resident macrophages may self-renew through local proliferation (Figure 2-figure supplement 1E) (86). In the regenerative PBS-control hearts, Mac 1, 4, and 5 expanded quickly at 1 dpci and gradually reduced back to steady state at 7 dpci, while Mac 2 and 3 expanded substantially over the first week post injury (Figure 3B). On the contrary, we noticed a dramatic reduction of Mac 2 and retention of Mac 1, 4, 5 and 6 in the non-regenerative CL-treated hearts over time after injury (Figure 3B). Lastly, the minor resident populations Mac 8 and 9 were diminished after cardiac injury and barely recovered in both conditions (Figure 3B). We then calculated the cluster contribution toward regenerative vs. non-regenerative conditions in a cell proportion analysis (Figure 3C and D). Coincidently, macrophage clusters enriched in regenerative conditions are the major resident clusters Mac 2 and 3, which were either dramatically decreased or barely recovered in CL-treated hearts (Figure 3C). Furthermore, low *ccr2* expression was found in these resident populations (Figure 2-figure supplement 3A), suggesting that they might have originated from embryonic-derived lineage, instead of circulatory/monocyte-derived lineage.

**Figure 3.**
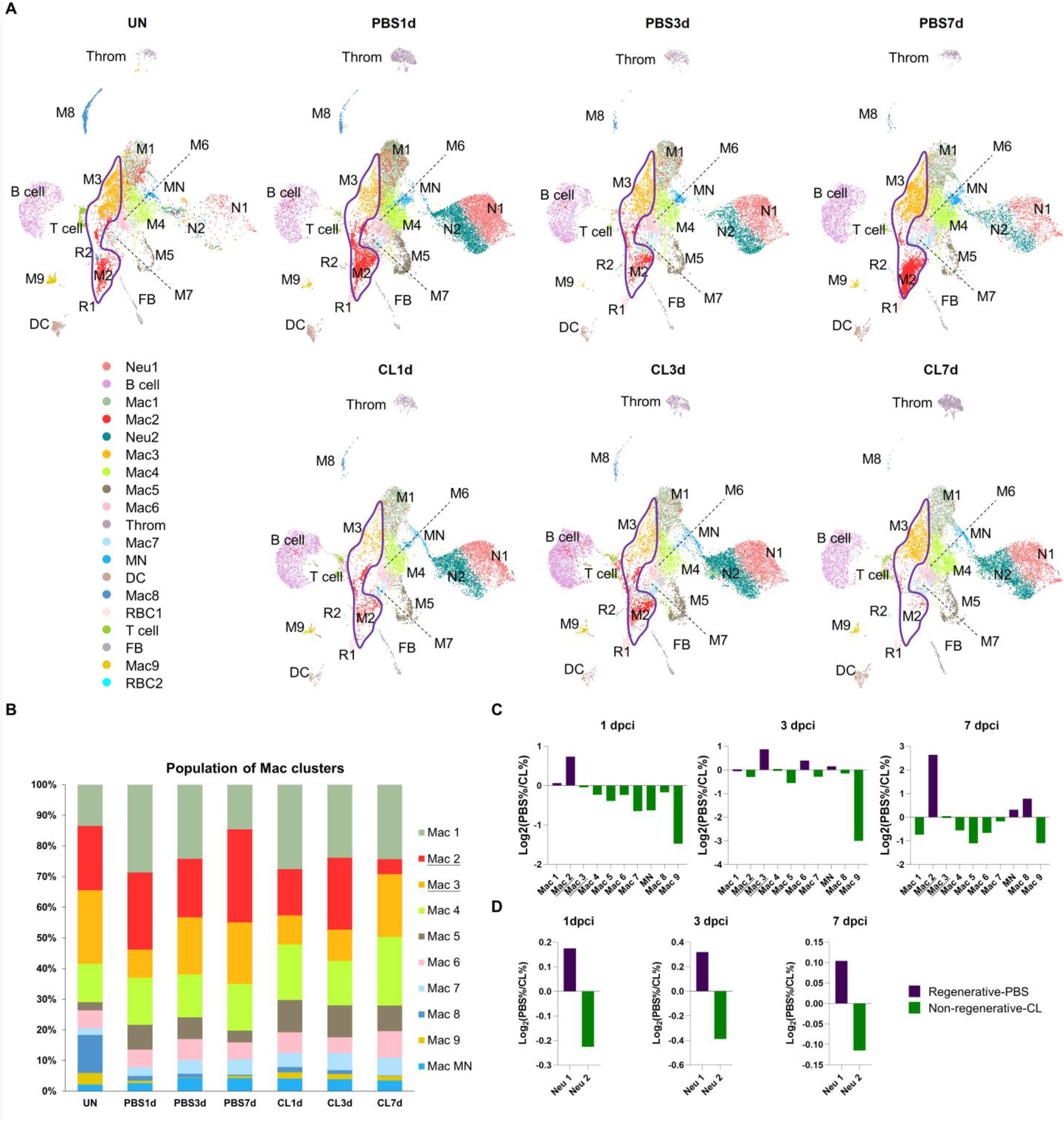
Temporal cell proportion analyses of inflammatory cell clusters identified resident macrophage clusters enriched in regenerative conditions. Differential proportion analyses of macrophage and neutrophil clusters under regenerative or non-regenerative conditions. (A) Split view of UMAP plots of major macrophage (Mac) and neutrophil (Neu) clusters as well as minor inflammatory cell clusters from uninjured and infarcted hearts under regenerative (PBS) or non-regenerative (CL) conditions. Mac 2 and Mac 3 clusters were the major resident macrophages enriched in regenerative conditions (delineated by purple lines) and they either dramatically decreased or barely recovered in non-regenerative conditions. (B) Stacked bar chart showed the percentage of macrophage clusters at each time point and condition. (C, D) Cell proportion analyses identified the regenerative-associated clusters (purple) and non-regenerative-associated clusters (green) of macrophages (C) and neutrophils (D). Proportion of each cell clusters under regenerative conditions vs. non-regenerative conditions were shown by log2 ratio.

On the other hand, Neu 1 and Neu 2 were two heterogeneous neutrophil clusters actively recruited to hearts after cardiac injury. We observed a decrease of both clusters from 3 to 7 dpci in the PBS-control hearts, while they retained in the CL-treated hearts by 7 dpci (Figure 3A). Interestingly, Neu 1 was increased in regenerative conditions, while the Neu 2 population was more prominent in non-regenerative conditions throughout the first week post cardiac injury (Figure 3D). These results delineate the dynamic changes of each inflammatory cell cluster and point to regeneration-associated resident macrophages, which were preferentially enriched in PBS-control hearts and might thus play pivotal roles in cardiac repair and regeneration.

### Differential function of inflammatory cell clusters in regenerative vs. non-regenerative conditions

To investigate the potential function of these heterogeneous inflammatory cells, we extracted and analyzed the overall differential expressed genes (DEGs) in each cluster (Figure 4A, **cluster-enriched DEGs**, full-listed in Figure 2-source data 2) and specific changes toward regenerative conditions (PBS-enriched) vs. non-regenerative conditions (CL-enriched) (Figure 4B, **condition-enriched DEGs**, full-listed in Figure 4-source data 1). Cluster-enriched DEGs reflect the general functions of each macrophage subpopulation, while condition-enriched DEGs reveal their property changes under regenerative vs. non-regenerative conditions, which may indicate their alternative activation. It has been shown that macrophages may originate from different niches and cell lineages, which can be roughly divided into embryonic-derived and monocyte-derived populations. These populations infiltrate and reside in each organ during embryonic development and after tissue injury (87). In addition, macrophages exhibit great plasticity and change their properties and functions in response to the microenvironment, which is often referred to as macrophage polarization (88). Both their origins and functional polarization may influence their gene expression profiles, so we perform differential expressed gene (DEG) analyses based on both cell clusters and regenerative vs. non-regenerative conditions to further depict their functions during cardiac repair and regeneration (Figure 4).

**Figure 4.**
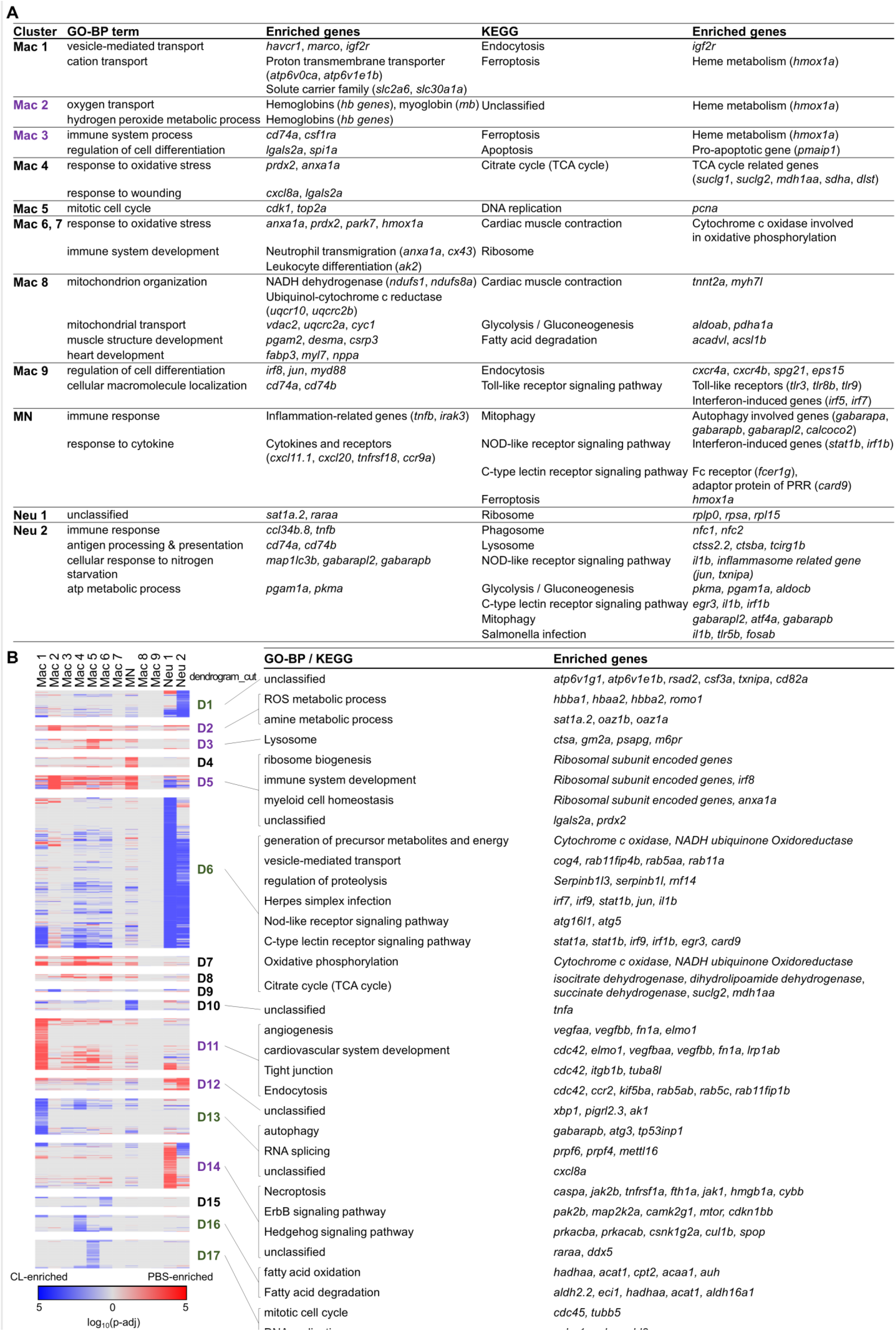
Differential gene expression in different inflammatory cell clusters under regenerative vs. non-regenerative conditions revealed alternative activation of both macrophages and neutrophils. Differential expression analyses of cluster-enriched and condition-enriched genes. (A) BP of GO and KEGG pathways were identified based on the cluster-enriched DEGs of each macrophage and neutrophil cluster. The major resident macrophage cluster Mac 2 and Mac 3 were highlighted in purple. (B) Hierarchical clustering of the condition-enriched DEGs between PBS and CL conditions of each macrophage and neutrophil cluster. PBS-enriched and CL-enriched genes were highlighted in red and blue, respectively. D1-D17 represented the dendrogram cut of the hierarchical clustering. BP of GO and KEGG pathways were identified from the DEGs in respective dendrograms. Regenerative and non-regenerative associated dendrograms were labeled by purple and green, respectively. DEGs, differentially expressed genes. DEGs, differentially expressed genes; GO, Gene Ontology; BP, biological process; KEGG, Kyoto Encyclopedia of Genes and Genomes.

Among regeneration-associated clusters (with higher cell proportions under regenerative condition in Figure 3), Mac 2 exhibited high expression levels of hemoglobin genes including *hbba1* and *hbaa1* (Figure 4A), as well as *romo1, prdx2* and *hmox1a* (Figure 4B; D2). Hemoglobin subunit expression was reported in murine macrophages upon LPS and interferon-γ stimulation (89), which may be involved in the transport of NO and regulate NO signaling and nitrosative stress (90, 91). *Romo1* functions as a redox sensor in mitochondria and regulate ROS production (92). *Prdx2* catalyzes the reduction of hydrogen peroxide and organic hydroperoxides to water and alcohols, which may serve a protective role against ROS stress upon cardiac injury (93). *hmox1a* is responsible for heme degradation and relative to iron homeostasis and inflammatory modulation (94). Preclinical studies suggest that heme oxygenase-1 (HO-1), encoded by *hmox1a*, degrades heme into cardioprotective and anti-inflammatory effectors such as biliverdin/bilirubin, carbon monoxide (CO), and free iron/ferritin (95). Biliverdin/bilirubin and CO further exert antioxidant and anti-inflammatory activities, upregulate genes such as *il10*, and promote macrophage polarization (94). These results suggest that Mac 2 may be involved in homeostasis including NO, ROS and heme regulation during the inflammatory and resolution phase, which was significantly diminished in CL-treated hearts at 7 dpci (Figure 3A and 3B). Mac 3 was enriched in genes involved in “immune system process” and “regulation of myeloid cell differentiation” including *cd74a, spi1a* and *lgals2a. Cd74* encodes the receptor for macrophage migration inhibitory factor (*mif*), known for its cardioprotective function in MI, suggesting a critical role for Mac 3 in innate immune response to cardiac injury (96). Mac 8 preferentially expressed CM structural genes such as *tnnt2a* and *myh7l*, and other genes involved in muscle structure and heart development, similar to the previously reported CX_3_CR_1_^+^ cardiac resident macrophages in mice (Figure 4A) (81). Mac 8 was also enriched in genes related to mitochondrial functions and energy metabolism, including both “Glycolysis/Gluconeogenesis” and “Fatty acid degradation”, involved in the metabolic switch in substrates preference for energy production (Figure 4A) (97, 98). It might reflect the roles of macrophage switching in pro-inflammatory and anti-inflammatory polarization at steady state or later repairing phase (99). Interestingly, Mac 8 was enriched in naïve hearts and barely recovered after injury in both conditions before 7 dpci (Figure 3A). It is intriguing to know whether cellular metabolism of this resident population relates to cardiac homeostasis and self-maintenance.

The macrophage clusters exhibited differential gene expression under regenerative vs. non-regenerative conditions, suggesting a functional polarization/alternative activation (Figure 4B). According to their expression profile dynamics, Mac 1 displayed diverse functions in response to cardiac injury under regenerative and non-regenerative conditions. Under regenerative condition, Mac 1 expressed genes related to the critical steps of regeneration—“angiogenesis” and “cardiovascular system development”, as well as debris clearance (*vegfaa, vegfbb, lrp1ab* and *elmo1*) (19, 100–102) and the regeneration-associated ECM protein-encoded gene, *fn1a* (Figure 4B; D11) (103). Under non-regenerative condition, Mac 1 expressed the inflammatory cytokines *il1b* and *tnfa*, as well as genes associated with “autophagy” and “mRNA splicing”, whose activity increase in response to inflammation (Figure 4B; D6, D10 and D13) (104, 105), suggesting an enhanced M1-like pro-inflammatory function. Furthermore, Mac 1 showed higher expression of *ccr2*, suggesting that this subpopulation might have derived from monocytes during cardiac injury, as described in murine cardiac macrophage classification (Figure 4B; D11) (5, 69). Among minor clusters, Mac 4, 6 and 7 showed higher expression of genes related to oxidative stress under regenerative condition, including *prdx2, anxa1a*, and *lgals2a* (Figure 4A and 4B; D5). *Anxa1* is a stress-induced gene which promotes neutrophil apoptosis, inflammatory resolution, and macrophage polarization during muscle regeneration (106, 107). Collectively, these three macrophage subpopulations might have roles in ROS homeostasis and in facilitating inflammation resolution. Under non-regenerative condition, Mac 4 expressed *cxcl8a* in response to wounding, a crucial chemokine for neutrophil recruitment in zebrafish (Figure 4; D13) (77), suggesting that Mac 4 may also play opposite roles under regenerative and non-regenerative conditions, coinciding with the enhanced recruitment and retention of neutrophils in CL-treated hearts. In addition, Mac 4-enriched genes were involved in “fatty acid oxidation” (*hadhaa, acat1*, and *cpt2*) and “TCA cycle” (*suclg1, suclg2*, and *mdh1aa*), but not in “oxidative phosphorylation” under non-regenerative condition (Figure 4A and B, D16 and D6). Incomplete fatty acid metabolism may cause TCA cycle intermediates accumulation (e.g., succinate), which may activate IL-1β and promote extensive ROS generation (108). Lastly, the subpopulations Mac 6 and Mac 7 were preferentially enriched in CL-treated hearts (Figure 3C) and expressed *anxa1a, cx43*, and *ak2* under both conditions which are related to neutrophil transmigration and leukocyte differentiation, suggesting that these macrophage subsets function in triggering neutrophil infiltration post cardiac injury (Figure 4A). Partly corresponding to the traditional concept of M1/M2 macrophage polarization, Mac 1, 4, and 5 preferentially expressed inflammatory genes, such as *il1b*, *tnfb*, and *ifngr2*, while Mac 2 and 3 preferentially expressed the M2 markers *arg2* and *mrc1b* (Figure 4-figure supplement 1A) (9, 78, 109–111). Taken together, detailed analyses on both clusters-enriched and condition-enriched genes suggest that each macrophage cluster might further exhibit distinct polarization states under different conditions, which reflect the complex nature of macrophage polarization and their potential roles in cardiac regeneration.

Besides macrophages, neutrophils are critical to tissue repair and regeneration (7, 112). The properties of zebrafish neutrophils are conserved with mammalian neutrophils in terms of development (113), morphology, and functions, including phagocytosis (114) and extracellular traps formation (115). In mice, infiltrated neutrophils contribute to cardiac healing by promoting macrophage polarization towards a reparative phenotype, while depleting neutrophils worsen the cardiac repair (116). However, neutrophil retention in damaged tissue leads to compromised tissue repair and regeneration by triggering a vigorous pro-inflammatory response (1). Among neutrophil clusters identified in our data, biphasic change was observed in gene expression under regenerative vs. non-regenerative conditions. Neu 1 was enriched in granule protein-encoding genes such as *lyz*, *npsn, mmp13a.1*, and *mmp9* (Figure 4-figure supplement 1B), while Neu 2 expressed inflammatory genes such as *il1b* and *tnfb*, phagosome-related genes *ncf1* and *ncf2*, as well as genes related to neutrophil chemotaxis such as *cxcr1, atf3*, and *illr4* (Figure 4A and Figure 4-figure supplement 1B). Although some expressed genes were observed in both neutrophil clusters, Neu 1 exhibited more up-regulated genes under regenerative condition (Figure 3D and 4B). For instance, high expression of retinoic acid receptor *raraa* indicates that Neu 1 might respond to retinoic acid produced in endocardium and epicardium after cardiac injury (Figure 4A and 4B; D14) (117, 118). Largely expressed DEAD-box RNA helicase *ddx5* (Figure 4B; D14) and spermidine/spermine N1-acetyltransferase 1 *sat1a.2* (Figure 4A and 4B; D2) were observed, indicating regulation of transcription and polyamine metabolism that associate with cell proliferation, differentiation, wound healing and tissue remodeling (119–121). The up-regulated gene *si:ch211-222l21.1* encodes prothymosin alpha that contains a protective role against ischemia-induced apoptosis (122), with enriched genes involved in necroptosis such as *fth1a, caspa*, and *hmgb1a* as well as genes related to inflammation regulation such as *cul1b* and *spop* (Figure 4B; D14) (123, 124). These results suggest that Neu 1 reacting to injury with functions related to debris degradation and clearance, inflammation regulation, and turn on programmed cell death in regenerative conditions. In contrast, Neu 2 was enriched in non-regenerative condition with increased expressions associated with inflammatory responses, such as *atp6v1g1* and *atp6v1e1b*, which encode for components of vacuolar ATPase that mediates vesicular acidification and contributes to pH related inflammatory responses (125). Moreover, *rsad2*, an interferon (IFN)-stimulated gene (126), was up-regulated (Figure 4B; D1). These enriched genes suggest that Neu 2 exhibits functions related to inflammatory propagation and recruitment of more inflammatory cells in non-regenerative condition (Figure 4A and Figure 4-figure supplement 1B) (127, 128). Both neutrophil clusters Neu 1 and Neu 2 expressed genes relative to energy metabolisms, regulation of proteolysis, and autophagy (Figure 4B; D6). These data indicate a change in energy homeostasis and differentiation in order to support prolonged survival of these neutrophils (129). Overall, these results explain to the continuous neutrophil recruitment and retention that we observed in the CL-treated heart and provide the potential molecular mechanisms underlying how inflammatory cells polarize during cardiac repair.

**Figure 4-figure supplement 1.**
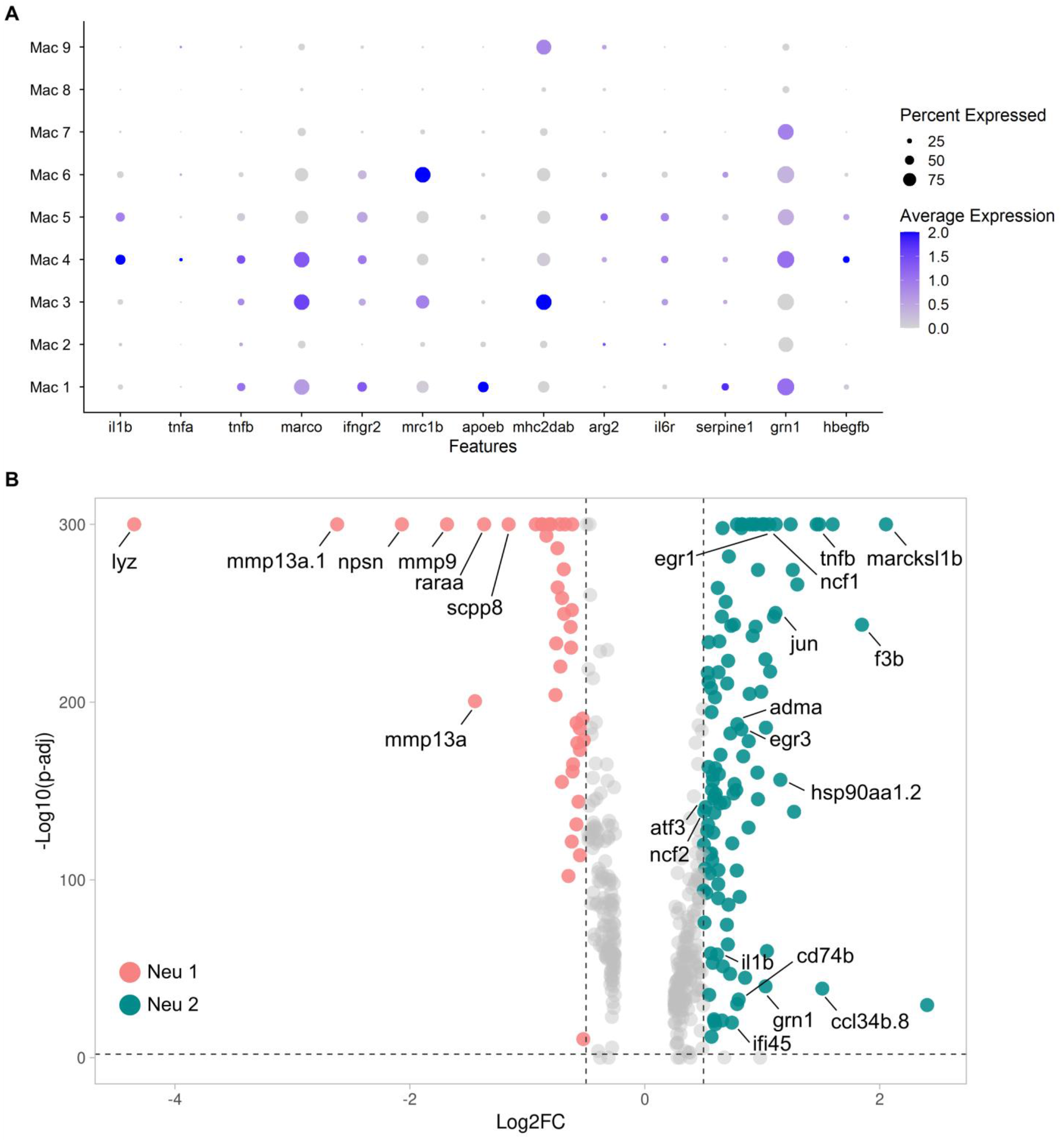
Expression pattern of Classical macrophage polarization genes and the DEGs between Neu 1 and Neu 2. (A) Dot plot showed the gene expression of common M1 and M2 markers across Mac 1 to Mac 9. The circle size represented the percentage of macrophages expressing the gene whereas the color represented the expression level. (B) Differentially enriched genes between Neu 1 and Neu 2 were shown in Volcano plot.

### Resident macrophages mediate ECM remodeling and phagocytic clearance of neutrophils by cellular crosstalk

Neutrophils are recruited to the injured tissue by various cytokines and chemokines and programmed for cell death as soon as they clear the tissue debris together with other professional phagocytes (130). Apoptotic neutrophils are cleared by macrophages to prevent further release of cytotoxic and inflammatory components, which is a critical step of inflammatory resolution (131). Our previous study and present results indicate that neutrophil retention occurs when macrophage properties are altered in non-regenerative condition, suggesting that the cellular interactions between macrophages and neutrophils may also change. This prompted us to investigate the cell-cell interactions between macrophages and neutrophils under regenerative and non-regenerative conditions. We first established the ligand-receptor pairs between macrophage and neutrophil clusters based on a published pipeline and then sorted them according to PBS-specific and CL-specific crosstalk at different time points post injury (Figure 5 and Figure 5-source data 1 to data 4) (132). It reveals that the putative interactions which change over time and under respective conditions.

**Figure 5.**
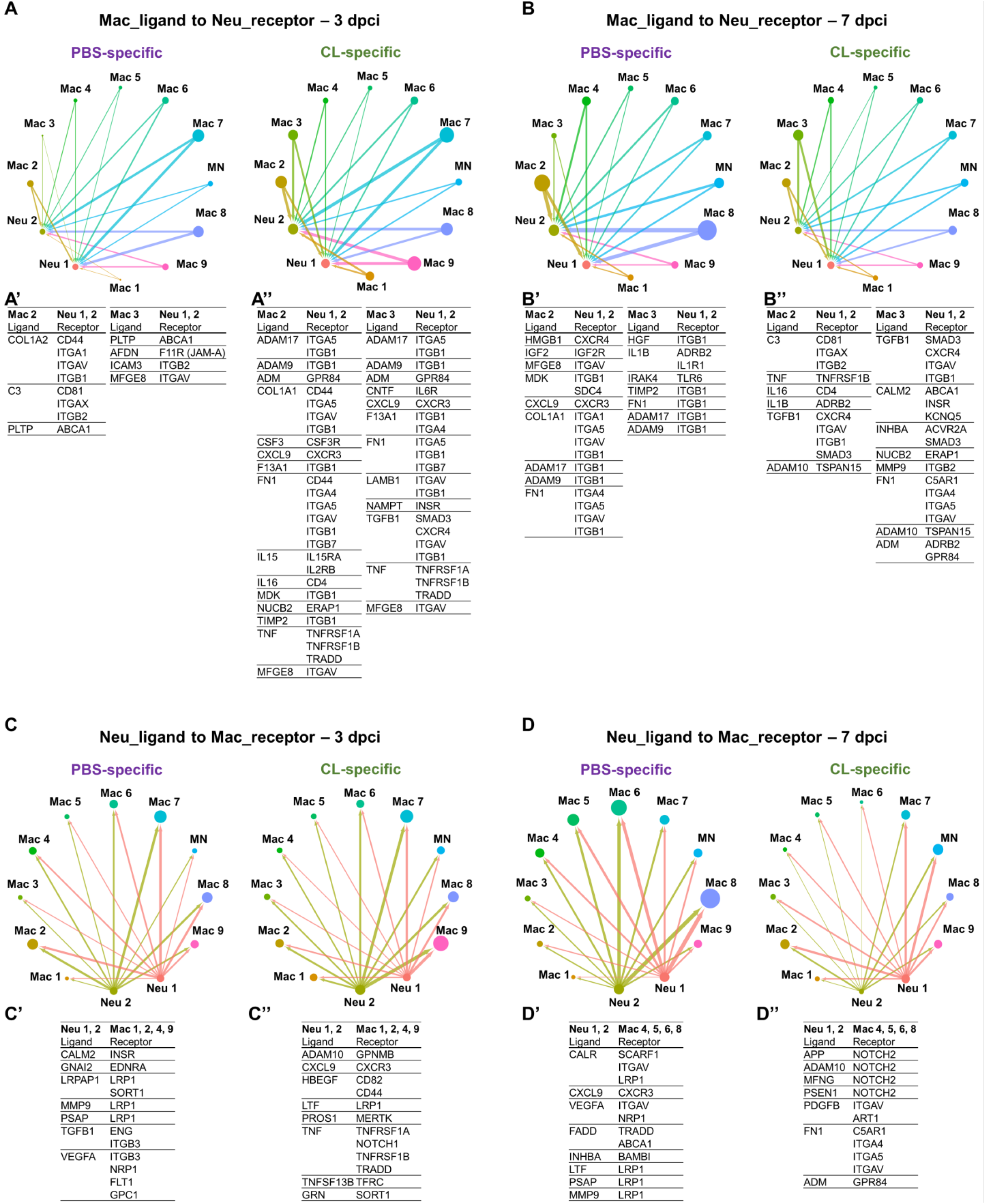
Cell-cell interactions between macrophages and neutrophils are altered in non-regenerative conditions. Crosstalk analyses identify hypothetical cell-cell interactions in macrophage and neutrophil clusters under regenerative (PBS) or non-regenerative (CL) conditions. (A and B) Putative interaction maps of macrophage-expressing ligands and neutrophil-expressing receptors between clusters at 3 dpci (A) and 7 dpci (B). Purple and green highlight the ligand-receptor pairs found specifically under PBS- or CL-treated conditions. Direction of arrows indicates the ligands signaling to the receptors in responding clusters. Circle size represents the numbers of ligand/receptor genes. Ligand-receptor pairs of the resident population Mac 2 and 3 to neutrophil clusters at 3 dpci (A’ and A”) and 7 dpci (B’ and B”) were shown. (C and D) Putative interaction maps of neutrophil-expressing ligands and macrophage-expressing receptors between clusters at 3 dpci (C) and 7 dpci (D). Purple and green highlight the ligand-receptor pairs found specifically under PBS- or CL-treated conditions. Direction of arrows indicates the ligands signaling to the receptors in responding clusters. Circle size represents the numbers of ligand/receptor genes. (C’ and C”) Ligand-receptor pairs of Neu 1, 2 to major macrophage responders at 3 dpci. (D’ and D’’) Ligand-receptor pairs of Neu 1, 2 to major macrophage responders at 7 dpci.

Among macrophage clusters, we focused on Mac 2 and 3 due to their enrichment under regenerative conditions (Figure 5A and 5B). Under regenerative condition, macrophage-ligand to neutrophil-receptor pairs involved in cell migration at 3 dpci, including the ligand complement component 3 (C3) with the receptor integrin ITGA1 (ITGAX/ITGB2), and afadin (AFDN) with the junctional adhesion molecule-A (JAM-A) (Figure 5A’) (133–136). Regulation of cholesterol homeostasis and inflammasome in neutrophils might also be regulated through PLTP–ABCA1 axis (Figure 5A’) (137, 138). At 7 dpci, adhesion, survival and migration of neutrophils were possibly regulated via ECM remodeling by macrophage expression of ADAMs, COL1A1, FN1, and TIMP2 (Figure 5B’) (139). Neutrophil chemotaxis might be provided by Mac 2 via the HMGB1–CXCR4 axis (140, 141) and neutrophil migration-related ligands MDK and CXCL9 (Figure 5B’) (142, 143). In addition, promotion of neutrophil self-phagocytosis could be mediated by the MFGE8–ITGAV axis under regenerative condition at 3 and 7 dpci (Figure 5A’ and 5B’) (144–146). On the other hand, more macrophage-ligand to neutrophil-receptor pairs were found under non-regenerative condition. At 3 dpci, more ECM components and regulators that could affect neutrophil behaviors were observed, including ADAMs, COL1A1, F13A1, FN1, TIMP2, LAMB1, and TGFB1 (Figure 5B”). To a lesser extent, the neutrophil migration-related ligands MDK and CXCL9 were also observed (Figure 5B”). Inflammation seemed to be further propagated through pro-inflammatory cytokine secretion from Mac 2 and 3, such as IL15, IL16, and TNF (Figure 5B”) (147, 148). Phagocytosis and survival of neutrophils could be positively modulated by IL15 (Figure 5B”) (149, 150). An increase of neutrophils was associated with CSF3 and ADM expression (Figure 5B”) (151, 152). In addition, neutrophil survival might be supported through the NAMPT-INSR and TGFB1-SMAD3 axes at both 3 and 7 dpci (Figure 5A” and 5B”) (153, 154), as well as the absence of clearance of apoptotic neutrophils via the MFGE8-ITGAV axis at 7 dpci (Figure 5B”). CXCR4 is involved in neutrophil reverse migration before apoptosis (155). The up-regulation of CXCR4 could be affected by the presence of TGFB1 at 3 and 7 dpci (Figure 5A” and B”) (156). These ligand-receptor pairs indicate differential neutrophil behavior resulted from altered macrophage properties and function, especially regarding the dynamic change in ECM remodeling, leading to enhanced neutrophil recruitment and/or retention under non-regenerative condition.

On the other hand, neutrophil-ligands to macrophages-receptors pair also showed dramatic differences between regenerative and non-regenerative conditions (Figure 5C and 5D). These ligand-receptor pairs were mainly involved in phagocytic clearance, when neutrophils express multiple eat-me/find-me signals recognized by macrophage receptors Lrp1 and Integrins, leading to neutrophil resolution only under regenerative condition. For example, neutrophils express calreticulin (CALR), a well-known “eat-me” signal, recognized by phagocytic receptors-LRP1 and SCARF1 on Mac 4, 5, 6, and 8 (Figure 5D’) (145, 157, 158). Furthermore, we observed various interactions mediated by the fas-associated death domain (FADD), FADD–TRADD and FADD–ABCA1 axes, which are related to the initiation of neutrophil apoptosis (Figure 5D’) (159, 160). By contrast, neutrophils in non-regenerative hearts expressed multiple ligands triggering NOTCH2 signaling which correlates with proinflammatory M1 macrophage polarization and the murine Ly6C^hi^ monocyte differentiation (Figure 5D”) (161, 162). Overall, the present crosstalk analyses indicate that neutrophils in non-regenerative hearts avoided resolution via programmed cell death and phagocytotic clearance, which led to their retention in hearts with chronic inflammation (inflammatory cytokines-receptors in Figure 5C”). These long-lived neutrophils may in turn prevent macrophages from polarizing towards pro-resolving/tissue-repairing phenotypes.

### Depletion of resident macrophage compromises heart regeneration

Based on the single-cell profiling results, regeneration-associated macrophages were mainly resident macrophages Mac 2 and Mac 3, which were substantially enriched in regenerative compared with non-regenerative hearts. To test the functional significance of these resident macrophages without disrupting the circulation/monocyte-derived macrophage recruitment, we perform CL depletion earlier at 8 days prior to cardiac injury (−8d_CL) (Figure 6). In our previous study, the effect of −1d_CL treatment only lasted a couple of days while the macrophage numbers soon recovered in adult zebrafish (1). We first examined the macrophage content in *Tg(mpeg1:mCherry)* reporter fish by flow cytometry at 2 and 8 days post CL injection (2 dpip and 8 dpip; Figure 6A). Indeed, the proportion of mCherry^+^ cells in the zebrafish hearts dropped from 1.7% at steady-state (green line) to 1.18% at 2 dpip (blue line), corresponding to the previous findings in −1d_CL treated hearts (Figure 6A) (1). At 8 dpip, the proportion of mCherry^+^ cells recovered to 1.64%, comparable to the steady-state level (Figure 6A, orange line). However, these recovered macrophages express *mpeg1*:mCherry at a lower level, which might reflect their differentiation status freshly from progenitors/monocytes (Figure 6A, orange line). These results also suggest that not all the resident *mpeg1*:mCherry^+^ cells are susceptible to CL depletion; for example, those B cells we observed from single-cell profiling (Figure 3A). Next, we performed cryoinjury on these fish after −8d_CL treatment and examined the macrophage content at 1 day post injury (1 dpci; Figure 6B). Surprisingly, the proportion of mCherry^+^ cells was even higher in −8d_CL group (1.14%) compared to PBS-controls (0.83%). Since scRNAseq results showed specific loss of Mac 2 and Mac 3, we examined their cluster-specific gene *hbaa1* and *timp4.3* expression by qPCR to test whether the Mac 2 and Mac 3 could recover from −8d_CL depletion (Figure 6C and Figure 6-figure supplement 1). Consistent with our scRNAseq results, the expression of both genes in *mpeg1*:mCherry^+^ cells were significantly reduced in −8d_CL hearts compared to PBS-controls (Figure 6D). These results support that early CL administration (−8d_CL) depleted resident macrophages without affecting overall macrophage infiltration, potentially derived from circulating monocyte or expanded from other resident clusters, after cardiac injury. However, Mac 2 and Mac 3 may not fully recover even when the overall macrophage numbers closely returned to the steady-state (Figure 6A, green line vs. orange line). Instead, they might be replaced by the monocyte-derived population with lower *mpeg1*:mCherry expression (Figure 6A). Of note, we again observed an overshoot of macrophage infiltration to the −8d_CL injured hearts, consistent with previously published results showing that −1d_CL treatment actually led to more macrophages infiltrated in the injured heart at 7 dpci (1). This observation indicates an intrinsic role of resident macrophages in modulating inflammation and immune cell recruitment after cardiac injury.

**Figure 6.**
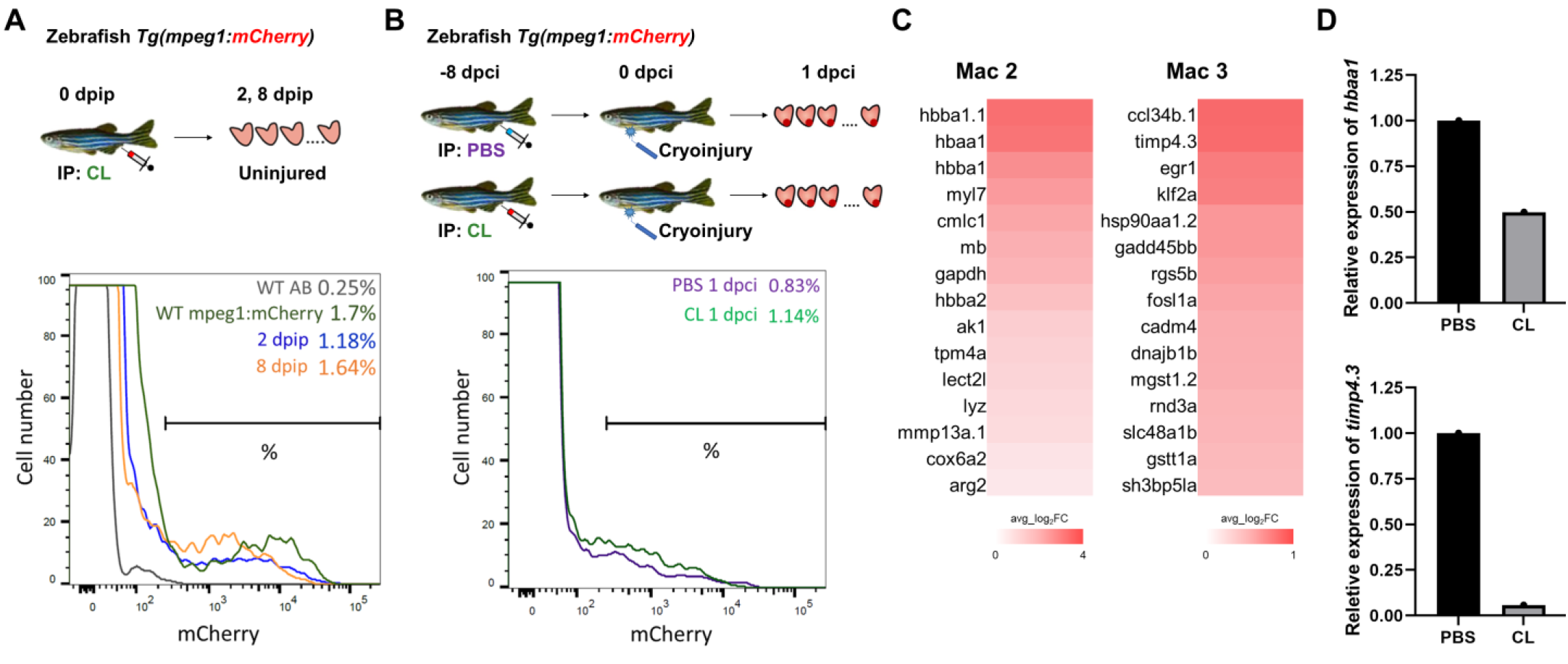
Early CL administration depleted non-recoverable cardiac resident macrophage clusters Mac 2 and 3. (A) Resident macrophage replenishment after CL depletion. Macrophages resided in the adult *Tg(mpeg1:mCherry)* fish hearts were examined by Flow cytometry at 2-dpip and 8-dpip after PBS vs. CL treatment. (B) *Tg(mpeg1:mCherry)* fish were IP injected with PBS or CL 8 days prior to cryoinjury. Cardiac macrophages (mCherry^+^) were examined by Flow cytometry at 1 dpci. (C) Heatmaps showed the top DEGs enriched in Mac 2 and Mac 3, respectively. The color represented the gene expression level in average log2 fold change. (D) Expression of Mac 2 enriched gene *hbaal* and Mac 3 enriched gene *timp4.3* in macrophages isolated from PBS- and CL-treated hearts at 8 dpip (without cryoinjury). The expression was examined by qPCR using *eeflalll* as the internal control. CL, Clodronate liposomes; IP, Intraperitoneal; dpip, day post intraperitoneal; dpci, day post cryoinjury; DEGs, differentially expressed genes.

To determine the functional requirement of resident macrophages in cardiac regeneration, we then examined the key processes underlying successful heart regeneration, including revascularization, CM proliferation and scar resolution in −8d_CL and PBS-control hearts at 7 dpci and 30 dpci (Figure 7A). Similar to our previous studies, we used *Tg(fli1:EGFP;myl7:DsRed-NLS)* fish to visualize the vascular endothelial cells in green and the nuclei of CMs in red, and used Acid Fuchsin-Orange G (AFOG) staining to reveal the scar size and composition in the injured hearts (1). Fast revascularization of the injured area in the first-week post injury is essential to support zebrafish heart regeneration (19). We examined revascularization by *ex-vivo* imaging of GFP^+^ endothelial cells in whole mount hearts and found that revascularization of the injured area in −8d_CL hearts were significantly decreased than PBS-controls at 7 dpci (Figure 7B and Figure 7-source data 1). We next examined CM proliferation by EdU incorporation and quantification on cryosection hearts, since dedifferentiation and proliferation of the existing CMs replenish the lost myocardial tissue from injury (163). Unlike previously reported in −1d_CL hearts, we did not observe a significant decrease in CM proliferation between −8d_CL and −8d_PBS hearts (Figure 7C and Figure 7-source data 1). Instead, we noticed small and round CM nuclei with weaker DsRed signals in the border zone of −8d_CL hearts compared to controls (Figure 7C). Correspondingly, the density of CM nuclei was significantly lower in the border zone of the −8d_CL hearts, suggesting that CMs survived less upon injury (Figure 7C). To test this possibility, we performed TUNEL assay to label damaged nuclei of apoptotic cells. Indeed, we found significant more TUNEL positive cells within the injured area of −8d_CL hearts (Figure 7D and Figure 7-source data 1). Coincidently, we also observed neutrophils retained in the injured area of −8d_CL hearts (Figure 7E and Figure 7-source data 1). As a result, −8d_CL hearts exhibited larger/unresolved scar tissues composed of both collagen and fibrin than −8d_PBS hearts at 30 dpci, reflecting compromised heart regeneration (Figure 7F and Figure 7-source data 1).

**Figure 6-figure supplement 1.**
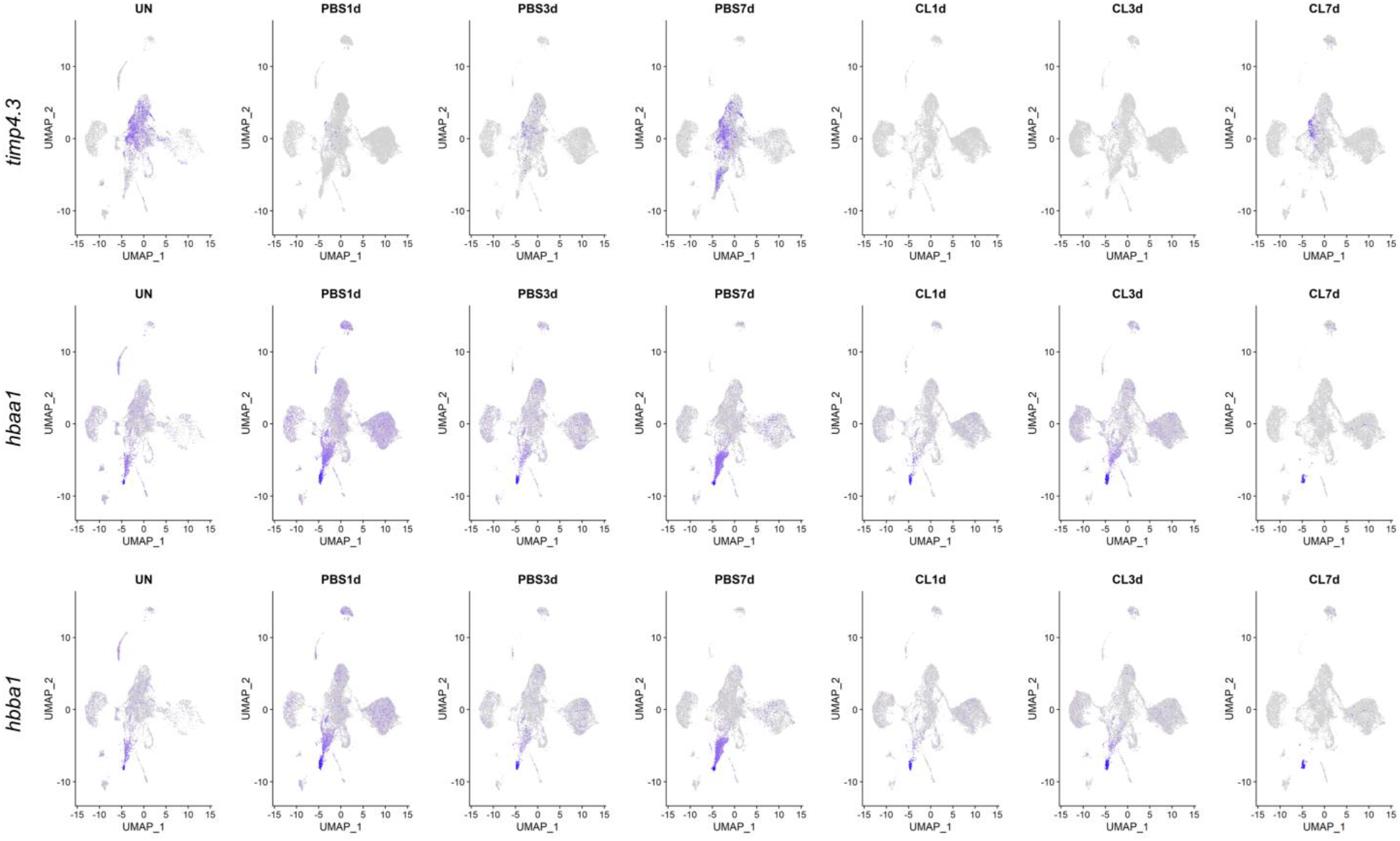
Expression of cluster-specific genes of resident macrophages Mac 2 and 3 in the regenerative vs. non-regenerative hearts post injury. Split UMAP plots for Mac 2 enriched genes *hbaa1* and *hbba1* as well as Mac 3 enriched gene *timp4.3* at each time point and condition. Both clusters and their marker genes remained diminished at 7 dpci. The color intensity represents the gene expression level.

**Figure 7.**
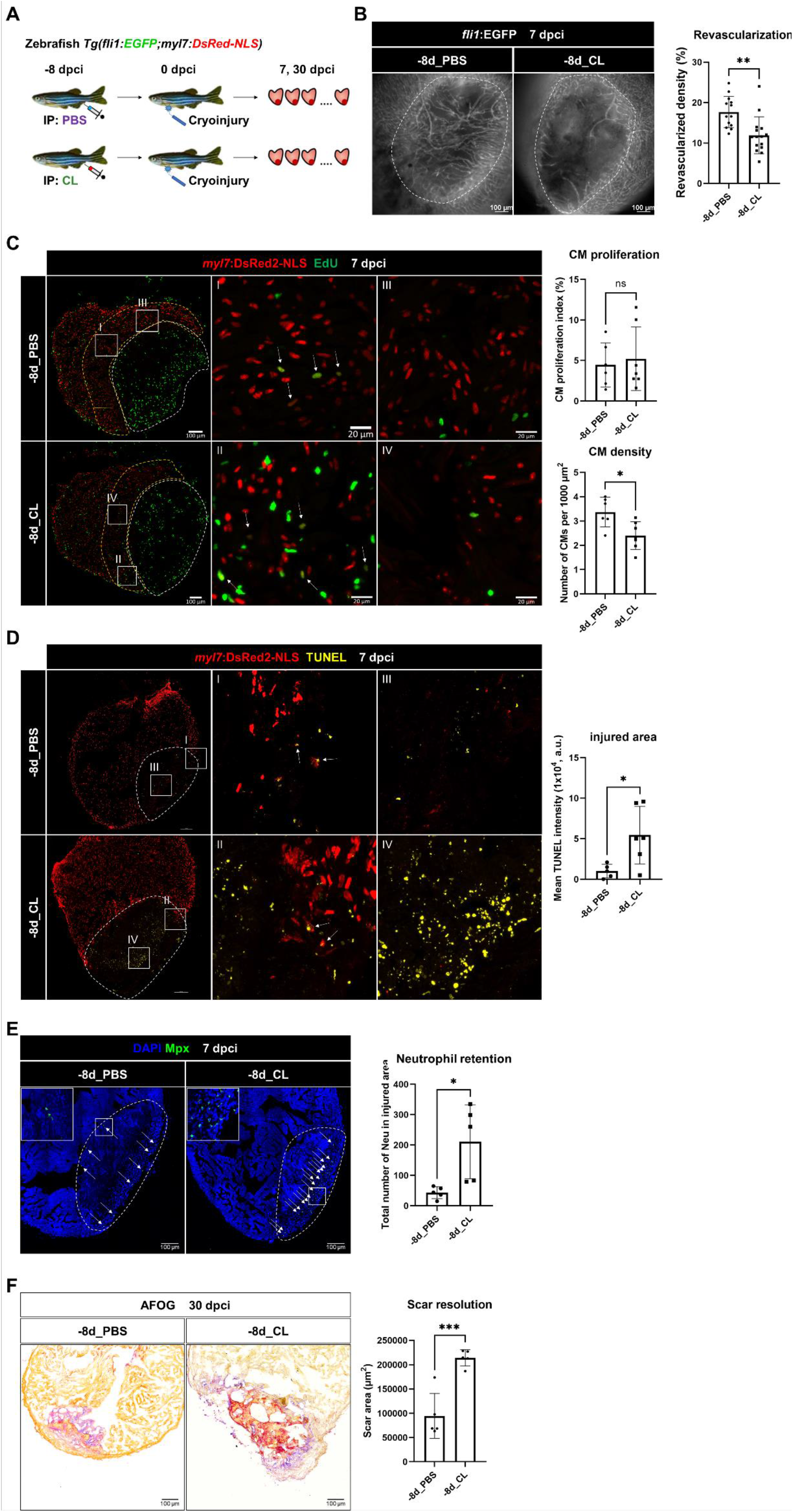
Depletion of resident macrophages compromised heart regeneration. (A) Experimental design for functional validation of resident macrophage depletion. Zebrafish were IP injected with PBS or CL at 8 days before cardiac injury (−8dCL). (B) Revascularization was evaluated at 7 dpci. Endogenous *fli1*:EGFP fluorescence depicted the vascular endothelial cells. White dotted lines delineated injury areas; scale bars, 100 μm. Quantification of vessel density by ImageJ was shown in the right panel. (C) CM proliferation was evaluated by EdU cell proliferation assay at 7 dpci. White dotted lines delineated injury areas; scale bars, 100 μm (left panels) and 20 μm (insets I–IV). White arrows pointed out the proliferating CMs (insets I and II). *myl7*:DsRed-NLS served as the endogenous CM nuclear marker. The shape of CMs became smaller and endogenous fluorescence was weaker in CL-pretreated hearts (−8d_PBS vs. −8d_CL, insets III vs. IV). Quantification of CM density and CM proliferation index in 200 μm adjacent to the injured area (delineated by yellow dotted lines) were shown in right panels. (D) TUNEL assay was performed on the same batches of cryosections which identified the CMs lost in the border zone (insets I and II) at 7 dpci. White arrows point out the TUNEL positive CMs in the area. More damaged nuclei were found in CL-treated hearts than in PBS controls (inset III and IV). Quantification of TUNEL intensity was listed in the right panel. (E) Neutrophils in injured areas were examined by Myeloperoxidase (Mpx) immunostaining. White dotted lines delineated injury areas; scale bars, 100 μm. Quantification of neutrophil number in injured areas was listed in the right panel. (F) Scar resolution was evaluated by Acid Fuchsin Orange G (AFOG) staining at 30 dpci. AFOG staining visualized healthy myocardium in orange, fibrin in red, and collagen in blue. Quantification of scar area was shown in the right panel. CL, Clodronate liposomes; IP, Intraperitoneal; CM, cardiomyocyte. The student’s t-test was used to assess all comparisons by Prism 9.

To test the potential ability of resident macrophages to recover after CL treatment, we performed CL administration 1 month prior to cryoinjury (−1m_CL, Figure 8 and Figure 8-source data 1). Strikingly, −1m_CL hearts still failed in regeneration, exhibiting significant defects in revascularization, neutrophil retention, and scar resolution (Figure 8). These results suggest that resident macrophages are essential for heart regeneration in modulating the revascularization, CM survival, and the resolution of inflammation and fibrotic scars, which cannot be replenished or functionally replaced by circulation/monocyte-derived macrophages once being depleted.

**Figure 8.**
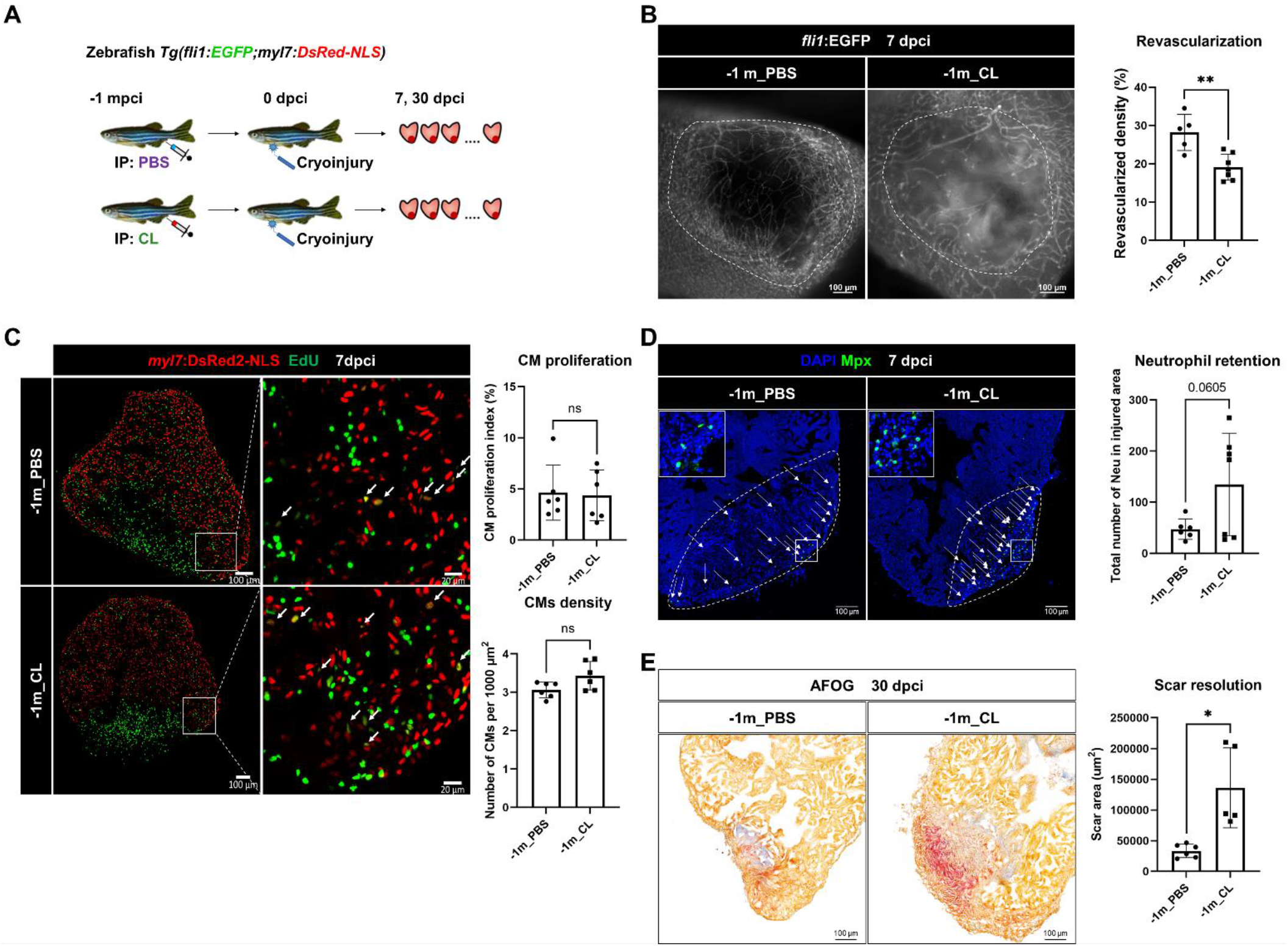
Depletion of resident macrophages led to long-term incompetence of heart regeneration. (A) Experimental design for functional validation of long-term depletion of resident macrophages at 1 month prior to cardiac injury. (B) Revascularization was evaluated at 7 dpci. Endogenous *fz’7*:EGFP fluorescence depicted the vascular endothelial cells. White dotted lines delineated injury areas; scale bars, 100 μm. Quantification of vessel density by ImageJ was shown in right panels. (C) CM proliferation was evaluated by EdU cell proliferation assay at 7 dpci. White arrows pointed out the proliferating CMs; scale bars, 100 μm (left panels) and 20 μm (middle panels). Quantification of CM density and CM proliferation index in 200 μm adjacent to the injured area was shown in the right panels. (D) Neutrophils in injured areas were examined by Myeloperoxidase (Mpx) immunostaining. White dotted lines delineated injury areas; scale bars, 100 μm. Quantification of neutrophil number in injured areas was listed in the right panel. (E) Scar resolution was evaluated by Acid Fuchsin Orange G (AFOG) staining at 30 dpci. AFOG visualized healthy myocardium in orange, fibrin in red, and collagen in blue. Quantification of scar area was shown in the right panel. CL, Clodronate liposomes; IP, Intraperitoneal; CM, cardiomyocyte. The student’s t-test was used to assess all comparisons by Prism 9.

## Discussion

### Timely inflammatory resolution and metabolic switch are critical events associated with macrophages function during zebrafish heart regeneration

Previously, Lai et al. showed that delaying macrophage recruitment in the first-week post cardiac injury by CL-mediated predepletion compromised zebrafish heart regeneration in terms of revascularization, neutrophil retention, CM proliferation and scar resolution, even though the macrophage numbers gradually recovered before 7 dpci. These previous data indicated a certain degree of functional divergence in the late infiltrating macrophages compared with the macrophages in the PBS-control hearts. Dynamics of inflammatory cells were also investigated during cardiac repair in mammals. Upon myocardial infarction, reperfusion is the standard practice in clinics, which salvages some CMs from ischemic death and thus reduce the infarcted area. In the mouse ischemic-reperfusion (IR) model, the vessel occlusion is released 30 mins after ligation to allow blood flow and immune cell trafficking, as compared to the permanent ligation (MI) model. Interestingly, inflammatory cell dynamic in the ischemic-reperfusion model resembles the regenerative zebrafish, when macrophage infiltration peak at an earlier time point, and neutrophils also resolve faster than in the permanent MI model, even though the cell numbers might not be comparable due to the differences in cell death and infarct size (164). Furthermore, macrophages from the IR model seem to repolarize and become inflammation-resolving M2 type faster than those from the permanent MI model, suggesting a progressive inflammation resolution (164). In the present study, we further characterized the PBS vs. CL models in zebrafish and found the transcriptomic changes upon macrophage delay. CL-treated hearts showed prolonged inflammation and dysregulated metabolism even until 21 dpci, suggesting that macrophages may play important roles in modulating neutrophil and inflammation resolution, which was further supported by the scRNAseq profiling of these inflammatory cells, discussed below. However, whether and how macrophages may directly regulate cardiac metabolism remain unclear. Metabolic shift in utilizing different metabolites for energy production is a critical process during cardiac repair and regeneration (34, 40, 41). Compared with PBS-control hearts, we observed downregulated genes predicted to be involved in both glycolysis and oxidative phosphorylation in CL-treated hearts from 7 to 21 dpci. The aberrant metabolism might be with macrophage phagocytosis function, since cardiac macrophages have been shown to preserve the metabolic stability of CMs by actively clearing the dysfunctional mitochondria and other waste via phagocytosis during homeostasis (43). Whether similar events occur during cardiac repair on top of their roles in debris clearance and how macrophages may directly influence CM dedifferentiating/proliferating await future investigation.

### ScRNAseq profiling revealed heterogeneous cardiac resident macrophages during steady-state and repair/regeneration

Since macrophages and neutrophils seem to be the main players in the regenerative PBS vs. non-regenerative CL conditions, we further profiled these inflammatory cells by scRNAseq. During cell isolation, we noticed a portion of *mpeg1*:mCherry^+^/*mpx*:EGFP^+^ cells in steady-state and enriched in PBS-treated hearts post injury. These double positive cells were confirmed as macrophages by cell sorting and Giemsa staining and were also described previously in the fin-fold amputated larvae (67, 165). From the scRNAseq profiling, we only observed a corresponding population of *mpeg1*^+^/*mpx*^+^ macrophages in Mac 2 at PBS7d. We speculate that EGFP proteins are stable in sorted *mpeg1*:mCherry^+^/*mpx*:EGFP^+^ macrophages and the *mpx* gene only needs to be turned on after injury and the replenishment of Mac 2 in PBS-treated heart. Myeloperoxidase-expressing macrophages were observed in human atherosclerotic plaque and increased during inflammation (166). Among macrophage clusters, Mac 1, 2, 3, 4, 9 are the major resident subsets at the steady-state (Figure 3B). The cluster enriched genes show similar functions to murine cardiac resident macrophages including phagocytosis/homeostasis, angiogenesis, antigen presentation, and sentinel function in monocytes and neutrophil chemotaxis (72, 87). However, the cluster/lineage markers were not comparable between fish and murine cardiac resident macrophages. The origin/lineage of fish cardiac macrophage is also difficult to determine without comparable markers and lineage tracing tools as in rodents. Therefore, new genetic tools are needed for further investigating these cardiac macrophage clusters. Nevertheless, judging by the dynamic change post injury and their replenishment in CL-treated hearts, Mac 2, 3, and 8 might be originated from the embryonic stage, as they show minor or no replenishment after depletion and injury. On the other hand, Mac 1 and 4 showed active recruitment to the injured heart and might be derived from monocytes (Figure 3B). After cardiac injury, the dynamic changes in numbers and gene expression among heterogeneous macrophage clusters reflect the swift response to injury and high plasticity of polarization, making it difficult to define regeneration-associated macrophage properties. Thus, we combined multiple analyses, including cell proportional analysis, GO analysis on cluster-enriched and condition-enriched DEGs, and crosstalk analysis in the present study. Interestingly, we observed some macrophage clusters other than Mac 2 and 3 turn on and off specific sets of genes under regenerative vs. non-regenerative conditions without affecting their overall identity (in clustering, Figure 4). This phenomenon suggests that functional polarization and dynamics might be associated with the presence of resident macrophages.

Interestingly, neutrophils were not only more in numbers, as they are retained and not cleared in the CL-treated hearts than those in the PBS-control hearts, but also exhibit different contributions of two subsets Neu 1 and Neu 2. Regeneration-associated Neu 1 showed enriched necroptosis gene expression but not classical neutrophilic genes involving inflammation, phagocytosis, and cell migration than non-regeneration-associated Neu 2. Further studies are required to functionally test the role of these neutrophils.

### Zebrafish cardiac resident macrophages are indispensable in heart regeneration

Our functional results reveal the essential roles of cardiac resident macrophages in zebrafish heart regeneration, as they are key to support revascularization, CM survival and replenishment, and eventually scar resolution. However, we did not observe cardiac resident macrophages directly contributing to CM dedifferentiation and proliferation. CM repopulation is affected possibly due to compromised revascularization (167), and the abnormal ECM deposition which leads scar deposition without affecting CM proliferation (103, 168). Moreover, ECM components provide various signaling events with other cell surface receptors that abnormal ECM dynamics could activate a range of disorders. For example, we observed resident macrophages-derived Mac 2 population interacting with neutrophils through ECM components such as fibronectin, ADAM metallopeptidases and tissue inhibitor of metalloproteinase 2 (TIMP-2) in regenerative conditions at 7 dcpi, whereas communication by non-regenerative Mac 2 mis-activated earlier (3-dpci) with distinct ECM components such as pro-fibrotic Factor XIII A chain (169). The present results indicate that ECM-associated proteins turnover and activities might contribute to establishing pro-regenerative microenvironment, as seen in murine cardiac repair and regeneration (170).

In summary, our study highlights the plastic and heterogeneous macrophages and neutrophils at steady-state and the first-week post injury. In addition, we molecularly and functionally characterized cardiac resident macrophages whose role is essential for heart regeneration. Further studies are required to investigate the role and function of distinct macrophage subsets and the molecular mechanism underlying how resident macrophages promote heart regeneration, which will help uncover why these functional properties are absent in adult murine cardiac resident macrophages. We anticipate these data will provide new insights for developing therapeutic strategies by engineering macrophages for cardiac repair.

## Acknowledgment

We thank core facilities at the Institute of Biomedical Sciences, including Light Microscopy, Pathology, and DNA Sequencing Core; Dr. Meiyeh Jade Lu and the High Throughput Genomics Core at Biodiversity Research Center for NGS work; the Innovative Instrument Project (AS-CFII-111-212) for cell sorting service at Academia Sinica. We also thank all members of the Lai group for their valuable suggestions and Dr. Michele Marass for the editorial consultancy. Research in Lai group has been funded by the Clinical Research Collaboration Grant from the Institute of Biomedical Sciences (IBMS-CRC108-P01), the Research Project Grant from the Ministry of Science and Technology (MOST 108-2320-B-001-032-MY2), and the Grand Challenge Project from the Academia Sinica (AS-GC-110-P7).

